# A dynamic metabolic flux analysis (DMFA) model for performance predictions of diverse CHO cell culture process modes and conditions

**DOI:** 10.64898/2026.01.11.698917

**Authors:** Jayanth Venkatarama Reddy, Nikola Malinov, Jason Souvaliotis, Eleftherios Terry Papoutsakis, Marianthi Ierapetritou

## Abstract

Bioreactor pH can significantly affect Chinese Hamster Ovary (CHO) cell metabolism, thus impacting glycoprotein titers. However, there is very limited literature on incorporating pH in mathematical models for CHO cell metabolism. To address this limitation, guided by recently published experimental data, we have curated a stoichiometric network and formulated phenotype-driven kinetic expressions to develop a Dynamic Metabolic Flux Analysis (DMFA) model. The DMFA model incorporates Critical Process Parameters (CPPs), notably bioreactor pH, basal and feed media nutrient composition, feeding times, and inoculation cell densities to predict bioreactor performance: cell growth rates, antibody titers, and nutrient and metabolite profiles. The DMFA model was trained on diverse fed-batch data of the CHO VRC01 cell line to regress the kinetic parameters. The model’s utility was demonstrated through experimentally validated model predictions of CHO-cell performance in intensified fed-batch cultures, perfusion cultures, and cultures with different media. Experimentally validated predictions of a culture with high initial cell density and increased feed addition (intensified fed-batch culture) showed that mAb titers similar to fed-batch culture can be achieved with shorter culture durations. Similarly, experimentally validated predictions of perfusion bioreactor performance showed that coupling historical fed-batch data with computational tools can be leveraged to predict continuous biomanufacturing performance. We thus demonstrate that the developed mathematical model can simulate culture performance outside of the training data set. This supports the predictive robustness of the framework and provides a valuable tool for bioprocess development of diverse culture modes.

**Highlights:**

- Experimentally measured fed-batch cell culture data was used to curate a reaction network. This reaction network was integrated with phenotypically driven kinetic expressions to yield a dynamic metabolic flux analysis (DMFA) model.
- The DMFA model can predict bioprocess performance indicators such as concentration of viable cells, mAb, amino acids, glucose, lactate, and ammonia.
- The model was developed to make these predictions under various process conditions such as bioreactor pH, media concentrations, feed supplementation schedule, and initial cell densities.
- Predicting and experimentally validating the impact of high initial cell density and increased feed media supplementation yielded in mAb titers similar to traditional fed-batch processes with much shorter culture durations.
- The application of the DMFA model trained on data from a traditional fed-batch process to predict perfusion bioreactor culture performance was successfully demonstrated and experimentally verified.
- The impact of AMBIC reference media on cell culture process performance was also predicted and experimentally validated. The predictions of amino acid metabolism yielded insights into improving the media.

## 1 Introduction

Therapeutic glycoproteins have emerged as significantly effective treatments for a variety of diseases due to their complex structures, which include post-translational modifications. Over the last few decades, Chinese Hamster Ovary (CHO) cells have emerged as the primary platform for producing monoclonal antibodies (mAbs) (Dhara et al., 2018). Development of a process to produce these therapies in CHO cells involves cell line development, media optimization, feeding schedule optimization, bioreactor parameter optimization, and scale up (Bielser et al., 2018). These processes are most often interlinked and require experimentation to identify the optimal operating conditions as a large number of critical process parameters (CPPs) (pH, temperature, media composition, feeding schedules, and seeding cell density) can influence bioreactor performance. For example, changes in bioreactor temperature or pH result in changes in nutrient uptake rates or inhibitor production rates (Ahn et al., 2008; Yoon et al., 2005). Hence, expensive and laborious experiments are required to screen many different media formulations, feeding schedules, and bioreactor conditions to maximize mAb titers. Production of therapeutic glycoproteins is moving away from the traditional fed-batch platforms to those of intensified fed-batch cultures and perfusion bioreactors. Intensified fed-batch bioreactors are inoculated at high cell densities (facilitated by N-1 perfusion bioreactors) to achieve similar antibody titers to fed-batch processes with much shorter culture durations by skipping the initial growth phase of traditional fed-batch cultures. Shorter run times are desirable as they can maximize the utilization of large-scale production facilities (Chen et al., 2018). Perfusion bioreactors utilize cell retention devices and continuous media exchange to achieve high cell densities (greater than 100 million cells/mL) and can operate for much longer durations while continuously producing mAb. At high cell specific productivities, perfusion bioreactors can lead to better process economics but also increased process complexity (Yang et al., 2019).

Reliable mathematical models can help streamline and optimize the manufacturing and process development-based research and development tasks by directing desirable experiments to achieve optimal process conditions (Chakrabarty et al., 2013; Sommeregger et al., 2017). Modeling of mammalian cell culture is complex as process performance can be impacted by concentrations of many metabolites (amino acids, sugars, vitamins, and inhibitors) and process conditions (pH, temperature, dissolved oxygen). Further complexities arise as the concentrations of these metabolites can change with culture duration in fed-batch processes. Kinetic models provide a simple empirical framework that relates nutrient concentrations to their uptake rates and cell growth rates via semi-empirically constructed ordinary differential equations. These models can be solved to resolve time profiles of nutrients and have been applied to determine feeding schedules (Kiparissides et al., 2015). Stoichiometric models are non-parametric and rely on metabolite uptake and secretion fluxes to make predictions of intracellular flux distributions. Combination of kinetic and stoichiometric models for mammalian cell metabolism can enable simulation of dynamic culture profiles while maintaining mechanistic fidelity (Reddy et al., 2023). Coupling kinetic and stoichiometric models provides a system that can be more dependable while making predictions outside of the original model training design space.

Published experimental data from our group has shown that bioreactor pH significantly affects CHO cell metabolism by impacting glucose uptake rates, lactate production rates, ammonia production rates, amino acid uptake rates, cell growth rates, and cell specific mAb production rates (Reddy et al., 2025). However, there are very few models (Ghodba et al., 2025; Hogiri et al., 2018) in the literature that explicitly include the effect of bioreactor pH while modeling CHO cell metabolism.

In this work, we have developed a dynamic metabolic flux analysis model that can predict temporal concentration profiles of glucose, lactate, ammonia, essential amino acids, nonessential amino acids, cell density, and mAb as a function of bioreactor pH, basal media composition, feed media composition, media supplementation strategy, and seeding cell density. This has been achieved by curating a reaction network (Section 3.1) and developing kinetic equations (Section 3.2) to model glycolysis, amino acid metabolism, and cellular growth rates. The model was trained (Section 3.4) on fed-batch bioreactor data generated for the CHO VRC01 cell line at pH setpoints of 6.75, 7.00, and 7.25 (Reddy et al., 2025). We used the DMFA model, trained only on fed-batch data to predict CHO cell performance in intensified fed-batch and perfusion cultures by changing the seeding cell density, bioreactor pH, feed media supplementation or process mode (perfusion). Subsequent intensified fed-batch and perfusion experiments validated the model predictions and demonstrated the DMFA model’s reliability. The model was further tested by predicting cell culture performance under a completely different media composition relative to the training data, namely the AMBIC reference media. Literature experimental data was used to validate predictions of process performance in the AMBIC reference media. (Cordova et al., 2023).

## 2 Cell culture

### 2.1 Cell line and culture media

CHO-K1 cell line (Clone A11 from the Vaccine Research Center at the National Institute of Health) stably expressing a broadly neutralizing anti-HIV monoclonal antibody (VRC01) was used in this study. CHO cells were grown in 125 mL Erlenmeyer flasks at 30 mL culture volume for passage and inoculum preparation. Erlenmeyer flasks used for inoculation were seeded at 0.4 million cells/mL and incubated at 37°C in an incubator configured to 85% relative humidity, 20% O_2_ and 5% CO_2_. HyClone ActiPro media (Cytiva, catalog number: SH31039.02) supplemented with 6 mM L-glutamine was used as the basal media. HyClone Cell Boost 7a (Cytiva, catalog number: SH31119.02) supplement and HyClone Cell Boost 7b (Cytiva, catalog number: SH31120.01) supplement were used as feed media.

### 2.2 Fed-batch bioreactor experiments

We used fed-batch experimental data from our lab as reported (Reddy et al., 2025). Briefly, fed-batch experiments were performed in 1 L Eppendorf BioFlo 120 bioreactors at 750 mL culture. The temperature of the bioreactor was controlled at 37°C. The setpoint for dissolved oxygen was 40% and agitation was set to 90 rpm by using a pitched blade impeller. Three different bioreactor pH values (6.75, 7, and 7.25) were studied by running biological triplicates for each condition. The inoculation seeding cell density in the reactors was 0.4 million cells/mL. The bioreactor pH was controlled by sparging with CO_2_ and pumping 6% sodium bicarbonate solution. A significant amount of sodium bicarbonate was added to the reactor for the pH 7.25 condition due to high lactate accumulation. The feeding protocol involved adding 22.5 mL of HyClone Cell Boost 7a and 2.25 mL of HyClone Cell Boost 7b daily from day 3 onwards. Starting day 5, glucose concentration was increased to 9 g/L every day by spiking 45% (w/v) sterile glucose solution (Sigma Aldrich, catalog number: 50-99-7). Antifoam C was added when foaming was observed. Samples were collected daily for measurements of cell counts, glucose concentration, lactate concentration, ammonia concentration, amino acid concentration, and mAb titer. Cell counts and viability were measured by using the trypan blue assay on the DeNovix automatic cell counter.

### 2.3 Intensified fed-batch experiments

Intensified fed-batch (high seeding cell density) experiments were conducted by inoculating CHO cells in 1 L Eppendorf BioFlo 120 bioreactors (750 mL culture volume) at a seeding density of 5 million cells/mL. To demonstrate the predictive capability of the model, we decided to operate the intensified fed-batch bioreactor at a pH of 7.12. The pH was controlled by sparging with CO_2_ and pumping 6% sodium bicarbonate solution when necessary. The dissolved oxygen was set to 40% and controlled by sparging with oxygen and air. The agitation was set to 90 rpm using a pitched blade impeller. The ActiPro basal media along with the HyClone 7a and 7b feed media was used. Glutamine was only added initially to bring its concentration to 6 mM. On day 1 of the culture, 22.5 mL of HyClone Cell Boost 7a media and 2.25 mL of HyClone Cell Boost 7b media were added to the reactor. From day 2 onwards, 38 mL of HyClone Cell Boost 7b media and 3.8 mL of HyClone Cell Boost 7b media were added to the bioreactor. On day 1, glucose was supplemented to the reactor to bring the glucose concentration to 5 g/L. From day 2 onwards, glucose was supplemented to the reactor to maintain a concentration of 9 g/L. Culture was stopped when viability dropped below 80%. Since this was used for model validation, only a single experiment was performed. Daily measurements of glucose, lactate, ammonia, amino acids, antibody titer, and cell density were collected.

### 2.4 Perfusion bioreactor experiments

We used metabolic data from perfusion bioreactor experiments in our lab as reported (Malinov et al., 2025). Briefly, perfusion experiments were performed in a 1 L Eppendorf BioFlo 320 bioreactor at a working volume below 1 L. For cell retention, a 20 cm long, 0.65 μm modified polyether sulfone (mPES) hollow fiber filter (Repligen, USA) was operated in tangential flow filtration (TFF) mode at a target shear rate of 1000 s^-1^ with a centrifugal pump (PuraLev® i30MU, Levitronix, Switzerland). The bioreactor was inoculated at a target seeding density of 0.4 million cells/mL in HyClone ActiPro basal media with 6 mM supplemental L-glutamine. Batch mode operation ensued until day three of culture, after which perfusion was initiated at a rate of 1 vessel volume per day (vvd^-1^). The perfusion media, perfusion rate control, and operating setpoints have been described previously (Malinov et al., 2025).Cell growth continued until a pre-determined viable cell density (VCD) setpoint, after which an intermittent cell bleed was initiated to maintain steady state. Daily samples were collected for offline measurements of VCD, % viability, glucose, lactate, ammonia, amino acids, and mAb concentration as described for the fed-batch experiments. During steady state operation, sample collection took place before and after the intermittent cell bleed.

### 2.5 Measurements of substrate, metabolite, and IgG concentrations

Measurements of glucose and lactate were performed using the YSI 2900 bioanalyser. Ammonia concentrations and osmolality were measured by using the BioProfile Flex bioanalyser (Nova). Amino acid concentrations were measured using an Agilent HPLC 1260 instrument. Amino acid standards (Agilent, catalog number: 5061) and a HPLC column (AdvanceBio Amino Acid Analysis column, Agilent, catalog number: 695975-322). A guard column was also purchased from Agilent (catalog number: 823750-946). The amino acids were derivatized with OPA for primary amino acids and FMOC for secondary amino acids as per the protocol provided by Agilent. Antibody titers were measured on the Agilent HPLC 1260 using protein A chromatography using a POROS A HPLC column (catalog number: 1502226; ThermoFisher). As per the protocol provided by ThermoFisher, mobile phase A consisted of 50 mM phosphate and 150 mM NaCl at pH 7.0. Mobile phase B consisted of 12 mM hydrochloric acid at pH 1.9. A UV detector (280 nm) was used to detect the mAbs. Additional method details are provided in the literature (Reddy et al., 2025). HPLC grade IgG standard (catalog number: MFCD00163923) was purchased from Sigma-Aldrich.

## 3 Model formulation

The goal of this work is to utilize mathematical modeling with historical fed-batch data to predict bioprocess performance in intensified fed-batch bioreactors, perfusion bioreactors, and under different media formulations. To achieve our goal, the following modeling criteria must be met.

- The impact of easily controllable bioprocess CPPs such as initial concentrations of nutrients (basal media formulation), initial viable cell density, bioreactor pH, and dynamic nutrient concentrations on bioprocess performance need to be modeled.
- The model must be capable of predicting dynamic process performance. Changes such as feed addition time and perfusion and bleed rates need to be successfully modeled.
- Bioprocess performance is inherently impacted by a large number of variables such as the concentrations of all amino acids, sugars, and metabolic inhibitors due to cellular requirements. Thus, these nuances need to be included in the model.

Kinetic models aim to predict dynamic or transient metabolic profiles by utilizing ordinary differential equations that depend on concentrations of metabolites and kinetic constants such as yield coefficients (Ben Yahia et al., 2015). Thus, providing an avenue to make time dependent bioprocess changes such as modulating feed supplementation time, temperature shift, or pH shift (Hogiri et al., 2018; Kiparissides et al., 2015). Application of these models to mammalian cell culture bioprocesses is challenging as many metabolic profiles (amino acids, inhibitors, sugars, cell density, and mAb titers) can influence bioprocess performance. Scaling these models to incorporate many metabolites leads to exponential increase in the kinetic constants. Estimation of these kinetic parameters and formulation of the kinetic equations requires extensive experimental data under diverse conditions. (Reddy et al., 2023).

The assimilation of mechanistic knowledge from genomic and isotope tracer data has led to the development of stoichiometric models for CHO cell metabolism (Park et al., 2024). These models are non-parametric and thus can be scaled to include many metabolites, but they cannot directly be used to predict transient metabolic profiles (Reddy et al., 2023).

Combining kinetic and stoichiometric models to form a dynamic metabolic flux analysis model or a dynamic flux balance analysis model can synergistically leverage the advantages of both constituents to yield a framework capable of predicting dynamic profiles for a large number of metabolites under various bioreactor process conditions if provided with CPPs as inputs. (Reddy et al., 2023). A dynamic metabolic flux analysis model has been previously reported (Nolan and Lee, 2011; Nolan and Lee, 2012). This model utilized 34 reactions and 24 metabolites in the stoichiometric reaction network and 12 kinetic equations to model dynamic profiles of glucose, lactate, viable cell density, mAb titers, ammonia, and 8 amino acids. The model also successfully predicted the metabolic switch of various metabolites such as lactate, ammonia, and alanine. Although this model achieved some of the goals mentioned above, there is room for improvement. All of the canonical amino acids can influence cell metabolism, thus depletion of amino acids not included in the literature model must be addressed/included in our modeling efforts. Bioreactor process parameters such as culture pH can significantly influence CHO cell metabolism and must be incorporated into the kinetic equations. The mass balance surrounding the dynamic metabolic flux analysis model must also be drafted to address fed-batch or perfusion culture modes.

All mathematical models rely on several assumptions that need to be understood prior to utilization of the model in different contexts. The assumptions underlying the dynamic metabolic flux analysis model are listed below.

- Stoichiometric models rely on the pseudo steady state assumption. The timescale of change of mammalian cell metabolism is fairly slow. For example, the doubling times for CHO cells used in this study can vary from 0.9 to 1.6 days. Hence, we assume that the solution from the stoichiometric model is constant over a short time horizon (0.1 days).
- Curation of the stoichiometric model: Stoichiometric models of CHO cell metabolism in the literature can vary in size. Some models consist of 34 reactions and 30 metabolites (Naderi et al., 2011), whereas others can consist of 6663 reactions and 2341 metabolites (Hefzi et al., 2016). The complexity of the stoichiometric model depends on the desired functionality. Curation of a stoichiometric model requires assumptions related to cell metabolism that have been derived from literature stoichiometric models and isotope labeling studies. This process has been described in Section 3.1.
- Kinetic equations are semi-empirical. This means that observed phenotypes and literature knowledge of cell metabolism are used to formulate the kinetic equations. The assumptions behind these formulations are described for each equation in Section 3.2.
- Lipids, vitamins, and trace metals can also significantly influence cell metabolism (Nargund et al., 2015; Stone et al., 2021). Eventually, models will need to include these contributions in some form. However, we assume that these metabolites have minor contributions to the experiments/predictions in this study as the concentrations of lipids, vitamins, and trace metals were not measured or modulated.

### 3.1 Metabolic-reaction network

We developed a reaction network consisting of reactions describing glycolysis, the TCA cycle, amino acid metabolism, the urea cycle, custom IgG formation, and biomass synthesis. We started with a reaction network from the literature consisting of 100 reactions and 72 metabolites (Zamorano et al., 2010). Certain metabolites (vitamins and unnecessary intermediate reactions) were not relevant for our applications, and their reactions were removed. Metabolic flux analysis (MFA) involves solving nonlinear optimization problems with linear constraints (described in Section 3.3). Constrained nonlinear optimization problems often involve finding the inverse of the stoichiometric matrix. To avoid computational challenges of matrix inversion arising from sources such as reaction scaling and collinearity, yet maintain enough relevant biological information, the reaction network has been formulated by lumping reactions and scaling stoichiometric coefficients to yield a low condition number (18) (Reddy et al., 2023). The process of lumping and scaling reactions is described in the supplementary section. This resulted in a reaction network consisting of 68 reactions and 42 metabolites shown in Figure 1 and supplementary Appendix Table A1. The IgG formation reaction was developed based on the amino acid sequence of the mAb used in this study. This reaction was scaled by dividing the stoichiometric coefficients of the amino acids with the total number of carbon atoms (6468) per molecule of mAb. Hence, the flux for the IgG reaction is also normalized by the same value while performing metabolic flux analysis. Biomass (CHO cells) reaction has been formulated based on CHO-K1 cell lipid, protein, RNA, and DNA composition reported in the literature (Hagrot et al., 2017; Széliová et al., 2020). CHO cell concentrations are measured in the units of million cells per mL. Since the biomass reaction has been formulated per Carbon-mol (Cmol) of biomass, we convert the viable cell density units to Cmols to generate the biomass flux in Cmols/cell/day for metabolic flux analysis. Hence, the biomass flux is calculated by utilizing 247 pg/cell and 24.12 g/Cmol of biomass (Berrios et al., 2011; Széliová et al., 2020). The process of deriving the biomass reaction is described in the Supplementary Section S1.

**Figure 1:**
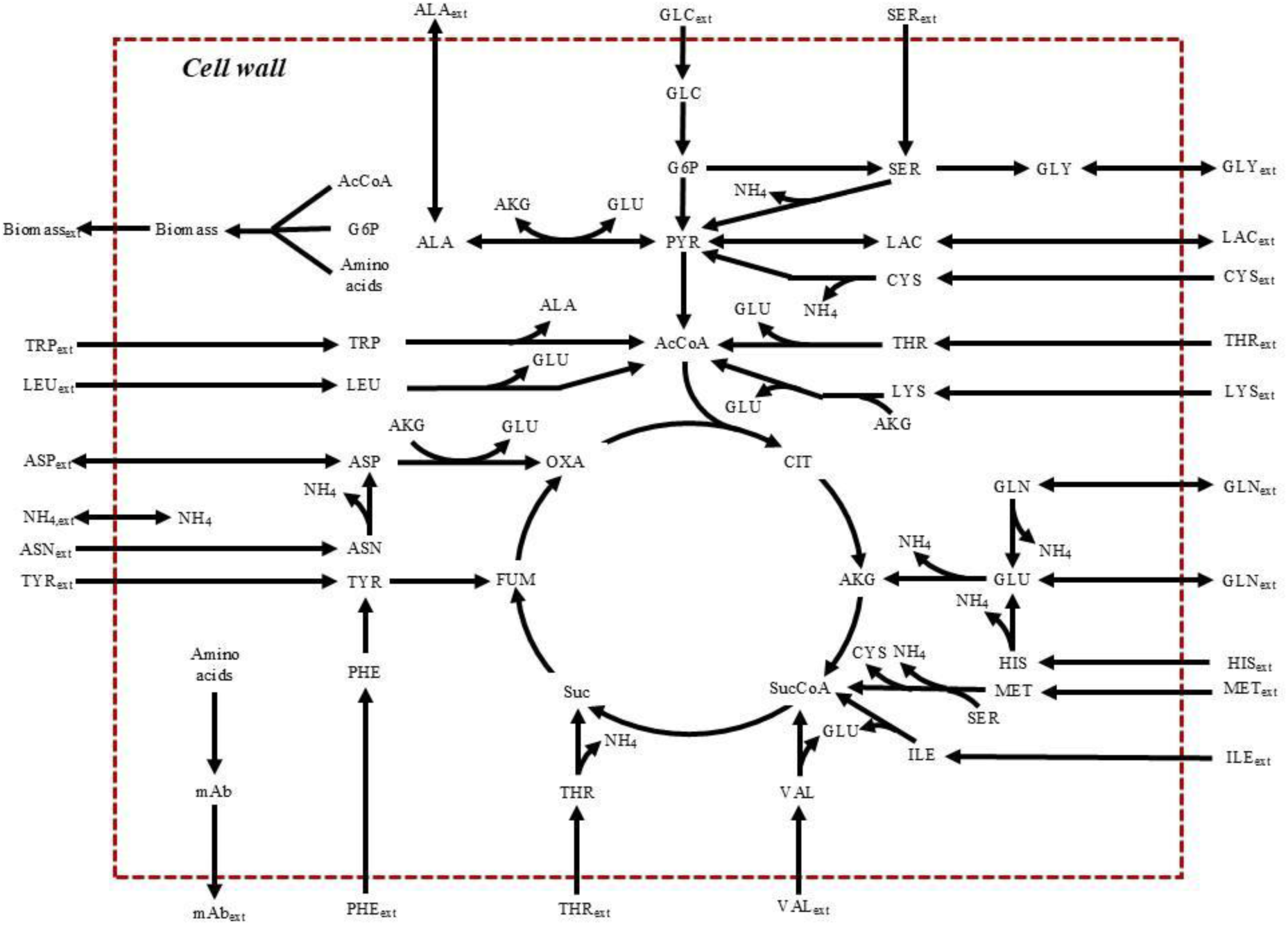
Schematic of reaction network used in the dynamic metabolic flux analysis model. Reactions with stoichiometric coefficients are listed in supplementary Appendix Table A1.

Our previous study applied a network consisting of 144 reactions and 101 metabolites (Reddy et al., 2025). While the network is compact, it is under constrained (more reactions than experimentally measurable fluxes) and solved through Flux Balance Analysis (FBA). FBA problems necessitate the careful selection of an assumed cellular objective function to address the solution degeneracy problem of under constrained networks (Reddy et al., 2023). The choice of objective function can significantly influence the resulting flux solution (Schinn et al., 2021). The goal in this work was to develop a dynamic model that can predict metabolic phenotypes across a broad range of culture conditions, which would inherently require the selection of different objective functions if using an under constrained network. We therefore curated the reduced network described above which is solved through MFA and does not require an assumed cellular objective.

### 3.2 Kinetic expressions

Dynamic metabolic flux analysis integrates stoichiometric modeling with kinetic models or systems of ordinary differential equations (ODEs). These kinetic equations are empirical in nature and various forms of kinetics have been used in the literature (Ben Yahia et al., 2021; Saa and Nielsen, 2017). We will be using kinetic equations of Monod type for the growth rates and yield coefficients for the metabolites. The yield coefficients is defined as the amount of product produced per amount of substrate consumed. For example, 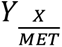 is the amount of CHO cells produced per amount of methionine consumed. Similarly, 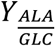 is the amount of alanine produced per amount of glucose consumed. The kinetic equations contribute to two key features of the model. The first is to correlate concentrations of metabolites to fluxes. The second is to make the framework dynamic. The integration of kinetic expressions into the dynamic metabolic flux analysis model to realize these key advantages is discussed in the subsequent section. Kinetic equations are semi-empirical, the assumptions and justifications behind framing the kinetic expressions are discussed in Appendix Table C1.

### 3.3 Dynamic metabolic flux analysis

The initial conditions to a dynamic metabolic flux analysis (DMFA) model are the seeding cell density, basal media concentration, feed media concentration, and feed media supplementation schedule. These initial conditions are fed to the kinetic equations to calculate the uptake rates of nutrients, production rates of byproducts, mAb production rate, and growth rate. The transport of these metabolites across the cell membrane are represented by reactions 46 to 68 in Appendix Table A1 and indexed by *j*. All the other reactions in the model are indexed by *i*. Rates (𝑣_𝑗,𝑚𝑒𝑎𝑠_) computed from kinetic equations are used to perform metabolic flux analysis as shown in Equation 1.

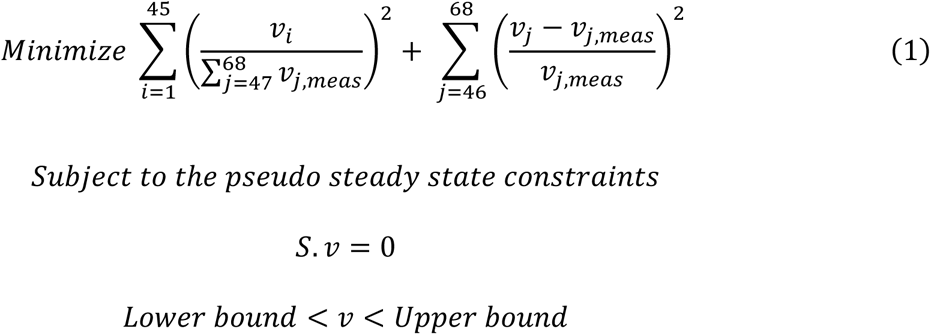

The objective function described in Equation 1 has two terms. The first term is used to minimize the sum of all the fluxes in the reaction network (except reactions 46 to 68). This term is normalized by the sum of all the kinetic expression predicted fluxes. Minimization of intracellular fluxes is based on the assumption that evolution with limited resources has placed pressure on living cells to operate with minimal effort (Holzhütter, 2004), this has led to the development of parsimonious enzyme usage flux balance analysis (pFBA) that has widely been used to provide more realistic solutions to stoichiometric models (Lewis et al., 2010). Minimizing the nutrient requirements after providing phenotypical constraints (constraining the biomass solution to the experimentally measured growth rate) has provided more accurate solutions to stoichiometric models as has been demonstrated for CHO cells in the literature (Chen et al., 2019). Hence, we have decided to include this term to minimize the intracellular fluxes while providing phenotypical constraints via a second term that minimizes the difference between the kinetic equation predicted flux and the stoichiometric model predicted flux. Thus, providing a solution that is closest to the kinetic equation predicted fluxes while satisfying biologically feasible constraints from the reaction network. Similar objective functions have been used in the literature to provide more realistic solutions to stoichiometric models (Jiménez Del Val et al., 2023). The matrix of stoichiometric coefficients (S) was constructed from the reaction network. The lower bounds and upper bounds were constrained to be 0 to ∞ if the reaction is irreversible and -∞ to ∞ if the reaction is reversible. Reactions 46 to 68 represent the phenotypical uptake and secretion rates modeled via kinetic equations. Hence, these values should be constrained to the fluxes provided by the kinetic equations. We constrain these fluxes by setting the lower bound and upper bound to 50% and 150% of the kinetic equation predicted flux. If no feasible solution is found at these bounds, the bounds are increased by increments of 10% and the stoichiometric model is solved till a feasible solution is found. The nonlinear optimization problem described in Equation 1 was solved by using IPOPT (Wächter and Biegler, 2006) implemented in python by using Pyomo (Hart et al., 2011) to determine all the intracellular fluxes (𝑣).

The results from the stoichiometric model were assumed to be constant for 0.1 days. The fluxes from the stoichiometric model and the concentrations of metabolites at time 𝑡 were used to calculate the concentrations of metabolites at time 𝑡 + 0.1. Details of these mass balance equations for fed-batch and perfusion bioreactors are available in Appendix Sections 10.1 and 10.2. The new concentration values (at time 𝑡 + 0.1 𝑑𝑎𝑦𝑠) were fed to the kinetic equations to determine the new fluxes at the next iteration point and the process was repeated. This process was repeated until harvest day or when viability dropped below 80%. A schematic of the dynamic metabolic flux analysis model is shown in Figure 2.

**Figure 2:**
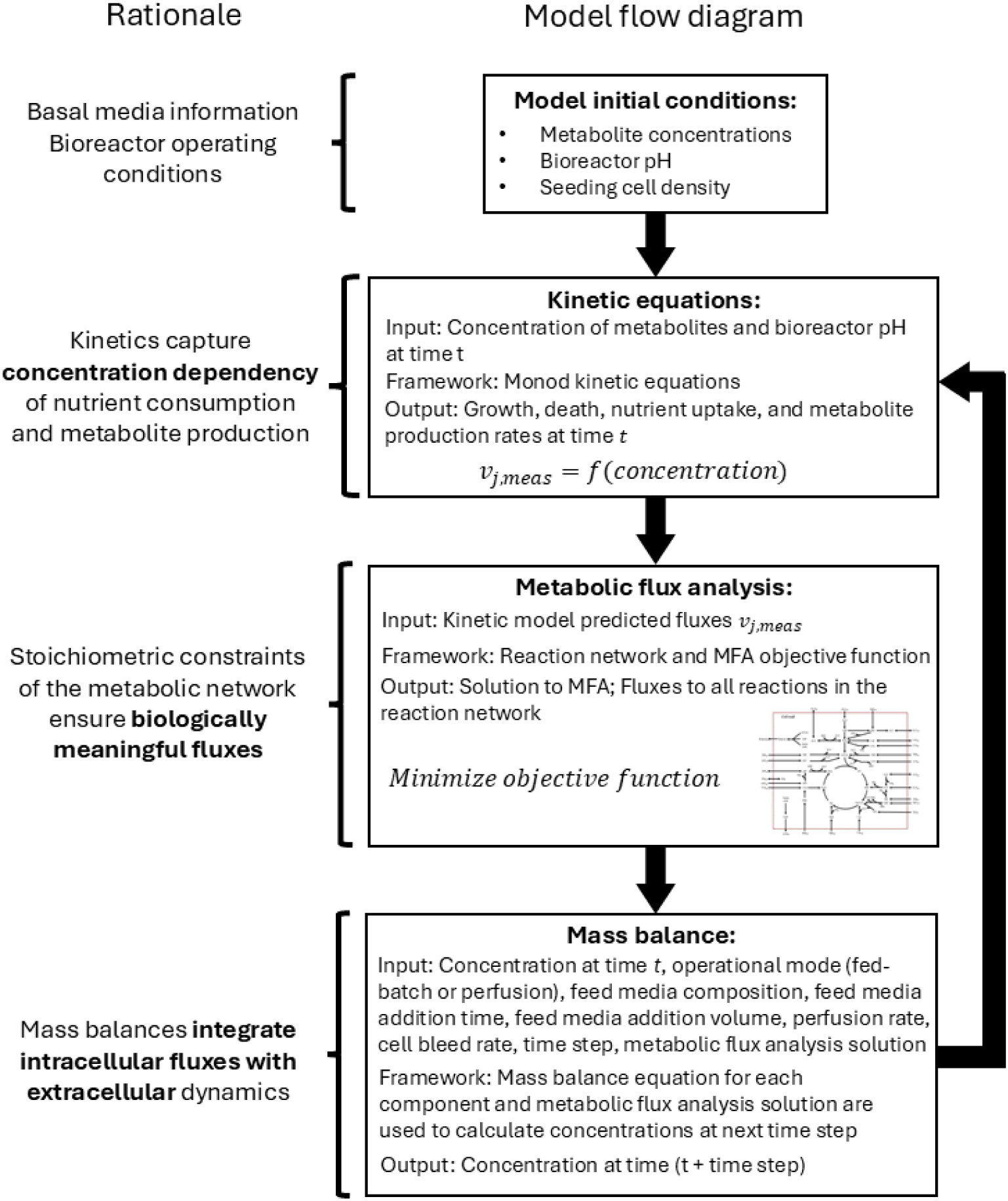
Schematic of all the steps in the dynamic metabolic flux analysis model along with the rationale for each step. The kinetic equations are provided in Appendix Table C1. The reaction network is available in Appendix Table A1. The mass balance equations are available in Appendix Sections 10.1 and 10.2.

### 3.4 Model regression

The DMFA model was trained on experimental data from experiments performed in our lab (Reddy et al., 2025) as described in Section 2.1.2. The measurements included 864 data points, the difference in the model predicted and the experimental value is represented by the sum of squared residual (SSR). Minimizing the SSR was used to determine the model parameters (𝜃) according to Equation 2.

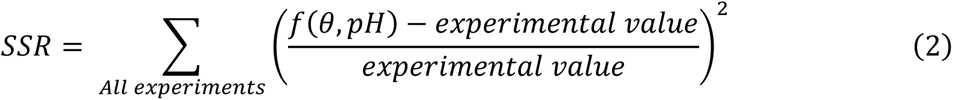

Equation 2 was minimized by using dual annealing (simulated annealing (Tsallis, 1988) with Nelder-Mead (Nelder and Mead, 1965)) using the python library SciPy (Virtanen et al., 2020). The results of regression are plotted in Figure 3. The model parameters are tabulated in Appendix Table B1 and B2.

**Figure 3:**
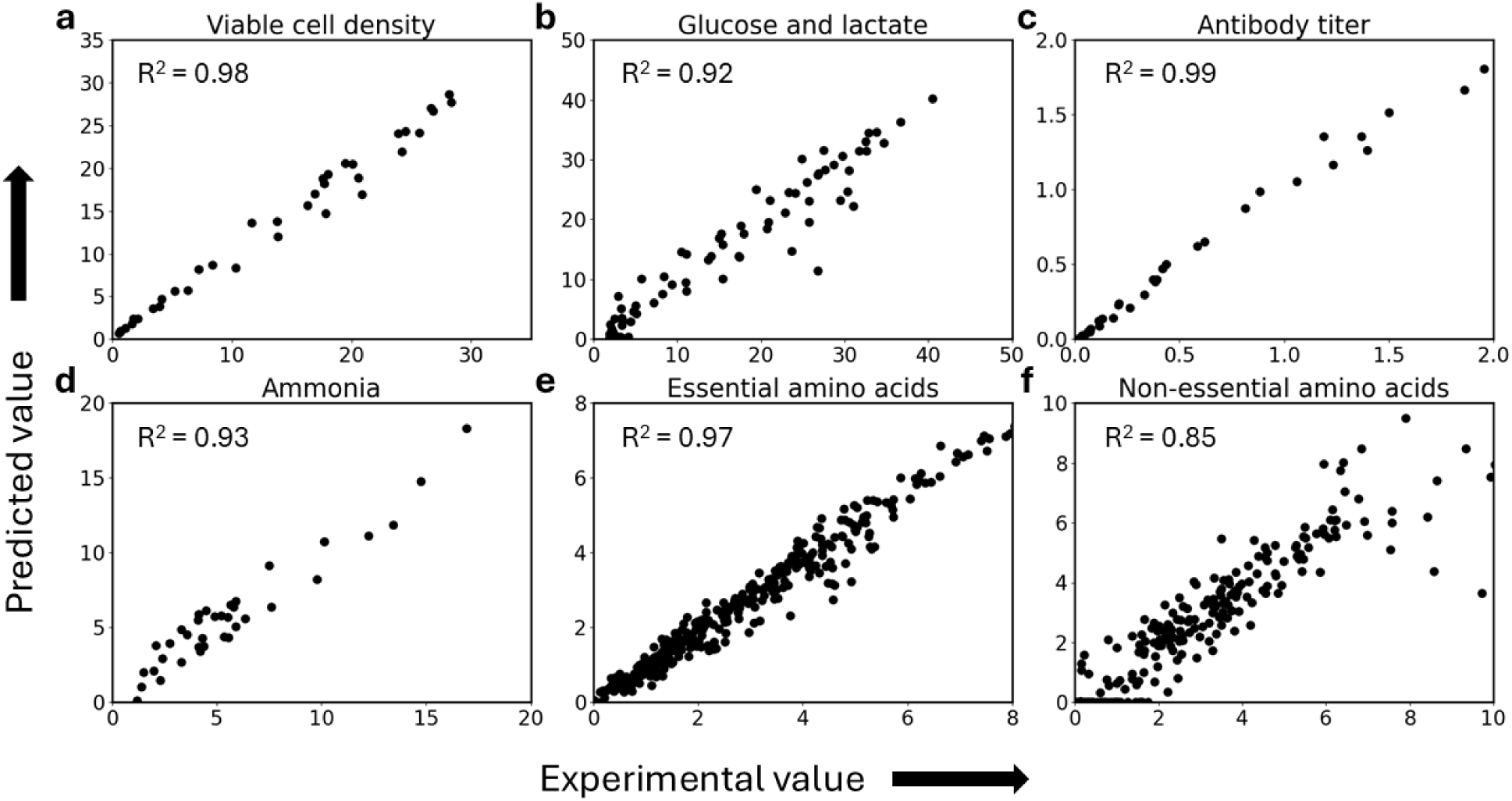
Scatter plot of experimental values vs the predicted values show that the model accurately captures variations in a) viable cell density, b) glucose and lactate concentrations, c) antibody titer, d) ammonia concentration, e) essential amino acid concentrations, and f) non-essential amino acid concentrations. The model predictions can describe variations in concentrations of ammonia, lactate, and non-essential amino acids but with slightly reduced accuracy.

High R^2^ values (greater than 0.95) were observed for regression related to viable cell density, mAb titers, and essential amino acid concentrations. Slightly lower values were observed for glucose, lactate, and ammonia concentrations due to modeling the complexities of lactate and ammonia metabolism. Lowest R^2^ values were observed for non-essential metabolism, also due to switch in metabolism of various non-essential amino acids. Nevertheless, the R^2^ values show that the model is capable of describing most of the variations in the experimental data. The spread of experimental measurements also highlights a very diverse dataset.

## 4 Results and Discussion

### 4.1 Impact of bioreactor pH on CHO cell metabolism

Equations C1 to C8 (Appendix C) describe the growth rate (𝜇) as a function of concentrations of glucose, glutamine, glutamate, asparagine, and aspartate. The cells had the highest growth rate from day 0 to day 3 for all bioreactor conditions. This growth rate is fueled by rapid glutamine consumption. According to the protocol to grow the CHO VRC01 cells, it is recommended to add glutamine only at the start of the culture (Cordova et al., 2023). Depletion of glutamine (Figure 4b) is also successfully modeled; this depletion leads to reduction in growth rates of the cells. Equation C8 shows that the cell can still consume glutamate to produce glutamine for their protein requirements. Hence, growth rates do not drop to zero after glutamine depletion and grow at reduced rates until the depletion of asparagine (Figure 4c). The concentration of asparagine is also successfully modeled throughout the culture duration for all the pH values (Figure 4c). Bioreactors operated at pH 6.75 had reduced growth rates and nutrient uptake rates, this led to accumulation of nutrients and increase in bioreactor osmolality. Increase in osmolality of CHO cell culture has shown to significantly reduce growth rates (Alhuthali et al., 2021). Hence, the osmolality term in Equation C7 accounts for reduced growth rates. The final phase of the culture saw reduced growth rates (due to nutrient depletion or osmolality) and cell death. CHO cell fed-batch cultures, eventually lead to cell death even if sufficient nutrients are supplied, this is because inefficiencies in cell metabolism can lead to production of many inhibitory metabolites (Kuang et al., 2021). Hence, integral viable cell density was used in Equation C25 to model cell death (Supplementary Figure S2). The predictions of cell death seem over estimated but the experimental measurement of dead cell density using trypan blue dye does not account for lysed cells (Klein et al., 2015).

**Figure 4:**
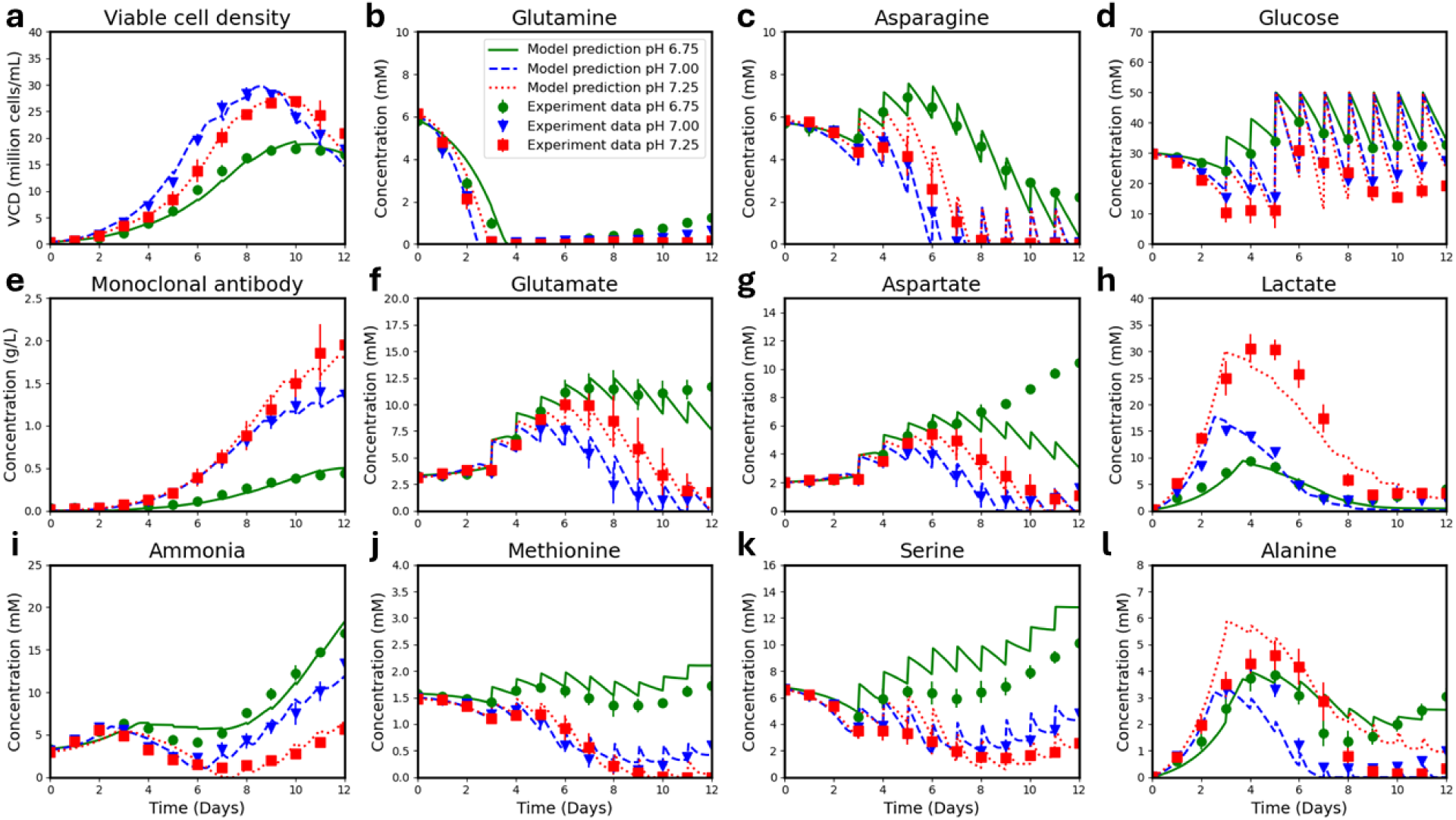
DMFA model can predict the effect of CPPs on growth rates, metabolite, and mAb concentrations in a fed-batch process. Successful regression of the model to fed-batch data is depicted by close alignment between experimental data and computational predictions for the concentrations of a) viable cells, b) glutamine, c) asparagine, d) glucose, e) mAb, f) glutamate, g) aspartate, h) lactate, i) ammonia, j) methionine, k) serine, and l) alanine.

The mAb concentrations were successfully modeled (Figure 4e) using a single parameter (𝑞_𝑚𝐴𝑏_) that is a quadratic function of bioreactor pH as shown in Equation C26. Bioreactor pH significantly impacted glucose and lactate concentrations. The glucose uptake rate was modeled by using two parameters, representing contribution of glucose towards growth 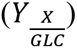 and the contribution of glucose to maintaining cellular metabolism (𝑚_𝐺𝐿𝐶_). This formulation successfully predicts the increase in glucose uptake rate with increase in bioreactor pH (Figure 4d) throughout the culture duration. Lactate production rates increased with increase in bioreactor pH leading to accumulation of lactate in bioreactor pH 7.25 condition. Lactate also exhibited a shift in metabolism from initial production to subsequent consumption. The production of lactate was modeled as a function of glucose uptake rates (Equation C10) with a yield coefficient 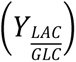 that is a function of bioreactor pH. The increase in glucose uptake rates and a higher value of the yield coefficient with increasing bioreactor pH led to successful prediction of lactate accumulation (Figure 4h). For this cell line, flux balance analysis has shown that glutaminolysis can lead to glycolysis flux being diverted to lactate (Reddy et al., 2025). Hence, the switch in metabolism of lactate was modeled as a function of glutamine concentration and according to Equation C11, depletion of glutamine leads to a switch in metabolism of lactate, this assumption led to successful predictions of lactate switch in all the conditions.

Uptake rates of essential amino acids were modeled using yield coefficients 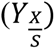 described in Equation C14. Bioreactors operated at pH 7.25 however, exhibited slightly increased amino acid uptake rates. Hence, the yield coefficients 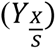 were scaled accordingly as described in Equation C14. A total of eleven yield coefficients and three quadratic constants for the amino acid scaling factor (𝐴𝐴𝑆𝑐𝑎𝑙𝑖𝑛𝑔(𝑝𝐻)) were used to model the concentrations of eleven amino acids (Figure 4j and Supplementary Figure S2). The increase in amino acid uptake rates at higher pH can lead to depletion of amino acids such as methionine (Figure 4j). This depletion has been successfully modeled by including a methionine term in the growth rate equation (Equation C8).

Asparagine (Figure 4c) and glutamine (Figure 4b) exhibited only consumption and were not produced throughout the culture duration. Equations C15 and C16 successfully predict asparagine and glutamine concentrations and their effect on cell growth. Consumption of glutamine led to production of glutamate and aspartate in the early phase of the culture. Glutamate and aspartate were consumed after glutamine depletion; this has successfully been modeled (Figure 4f and 4g) by Equations C17 to C20. Alanine was also produced during the early phases of the culture. Since it is also a byproduct of glycolysis, excess pyruvate accumulation can also lead to alanine production during the early phases of the culture. Hence, its predictions are very similar to the lactate profiles and Equations C12 and C13 can successfully predict alanine (Figure 4l) concentrations. Prediction of glycine metabolism was challenging for the model to describe as it appears that Equation C21 perform poorly at certain points of the culture leading to reduced R^2^ values (Supplementary Figure S2). Hence, caution should be exercised to utilize this model for predictions of glycine concentrations. Glycine metabolism appears only partially correlated to glutamine concentration and improved mechanistic understanding for the reason for the switch in metabolism or the utilization of statistical modelling approaches to generate hybrid models might be useful.

Ammonia metabolism is also extremely complex given the many sources and sinks for ammonia. The effect of glutamine consumption on ammonia metabolism, the subsequent consumption of ammonia, and the switch again from consumption to production has been successfully modeled using Equation C23. However, the R^2^ value of 0.93 (Figure 4i) indicates that ammonia predictions are not always accurate.

### 4.2 Prediction of intensified fed-batch cultures

Model validation requires experimental data that were not used to train the model. In addition to validation, we specifically wanted to demonstrate that our model is capable of predicting intensified fed-batch culture performance after training on only fed-batch data. Utilizing N-1 perfusion to seed the bioreactor at a higher cell density has led to the development of intensified fed-batch platforms for mAb production (Chen et al., 2018). Thus, demonstrating industrially relevant model applications such as derisking of intensified fed-batch process development. We collected intensified fed-batch data at a different bioreactor pH (7.12), an over 10-fold higher seeding cell density (5 million cells/mL), and a new feed supplementation plan (discussed in Section 2.3) that led to increased nutrient feeding.

Concentrations of metabolites that impact growth rates (glutamine and asparagine) were successfully modeled in the intensified fed-batch cultures (Figure 5b and 5c). This led to successful prediction of cell densities at the early cell growth phase (driven by glutamine), late cell growth phase (driven by asparagine), and cell death phase in the intensified fed-batch experiment at a pH 7.12 (Figure 5a). Bioreactor pH significantly impacted glucose uptake rates (Figure 4d). The predictive abilities of the model are demonstrated via successful glucose predictions (Figure 5d) throughout the culture duration in the intensified fed-batch process. Successful predictions of glutamine concentration (Figure 5b), led to a successful prediction of an early shift in lactate (Figure 5h), alanine (Figure 5l), and glutamate metabolism (Figure 5f). The increase in lactate accumulation at the new pH setpoint and subsequent depletion was also successfully modeled. The concentration of alanine was also successfully captured (Figure 5l) throughout the culture duration. The R^2^ square value for essential amino acid prediction is very high, hence it is not surprising that essential amino acid concentrations can be successfully modeled (Figure 5 and Supplementary Figure S3). This highlights the capability of the model to derisk the process prior to experimentation. The experimentally validated model predictions show that phenylalanine (Figure 4j) and threonine (Figure 4g) are accumulating in the culture. The experimentally validated model predictions also show that methionine is depleted in the culture (Figure 4k). Hence, indicating the need to rebalance the media formulation to provide appropriate nutrient levels to prevent depletion or accumulation. For example, experimentally validated model predictions show that histidine is supplied at the perfect level to maintain a constant concentration in the culture (Figure 4i). The predictions of other amino acids and ammonia are shown in Supplementary Figure S3. The model could successfully predict ammonia concentrations in the early phase of the culture but performed poorly in the later stages (Supplementary Figure S3). The most important experimentally validated model prediction is the mAb concentration (Figure 4e). This shows that the intensified fed-batch cultures can achieve similar mAb titers (on day 8) while compared to the fed-batch culture (on day 12) reducing the operational time for large scale bioreactors.

**Figure 5:**
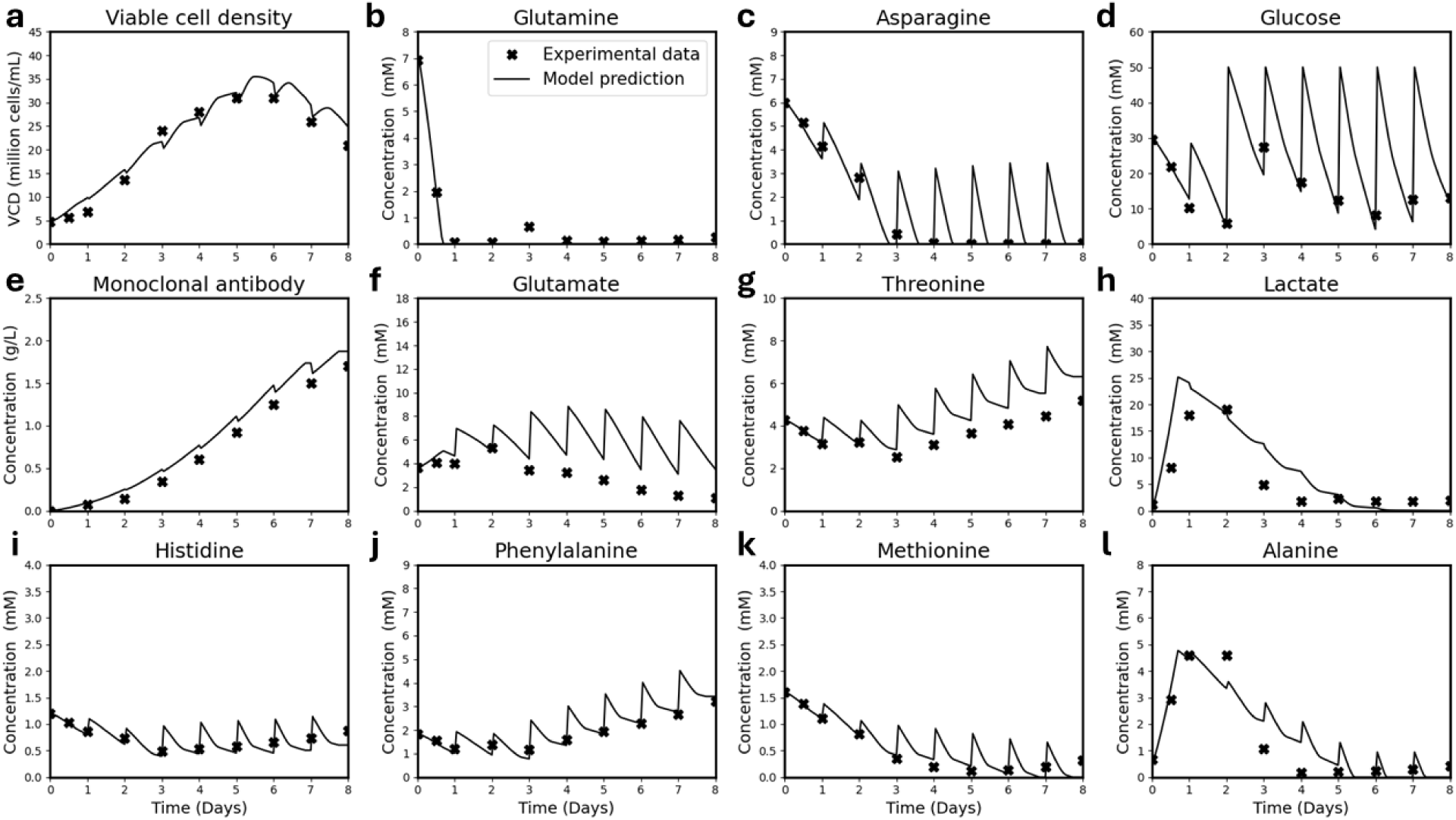
The model successfully predicts intensified fed-batch culture performance for concentrations of a) viable cell density, b) glutamine, c) asparagine, d) glucose, e) monoclonal antibody, f) glutamate, g) threonine, h) lactate, i) histidine, j) phenylalanine, k) methionine, and l) alanine.

### 4.3 Prediction of perfusion bioreactor cultures

In this section we utilize the DMFA model (trained only on fed-batch data) to predict performance in perfusion cultures. Figure 6b shows that the model accurately predicts glutamine depletion in the culture during the initial three-day batch period. Subsequent growth after perfusion initiation is driven by glutamate consumption that is also accurately predicted (Figure 6f). A scheduled intermittent cell bleed was used in the perfusion experiment as described in Section 2.4. Hence, the predictions of viable cell density drop whenever the cell bleed occurs at steady state. The viable cell density in perfusion with a target setpoint of 20 million cells/mL has been successfully predicted across the initial batch, transient, and steady stage regimes (Figure 6a). The model also successfully captures the metabolic switches of lactate (Figure 6h) and alanine (Figure 6l) during the batch and transient periods as the VCD increases, and their subsequent depletion or near depletion at steady state. The residual steady state concentrations of glucose and all the other amino acids have also been accurately captured by the DMFA model. The residual mAb concentrations (Figure 6e) has also been successfully predicted. Hence, we demonstrate the model’s ability to aid perfusion process development using only historical fed-batch data for simulation prior to experimentation.

**Figure 6:**
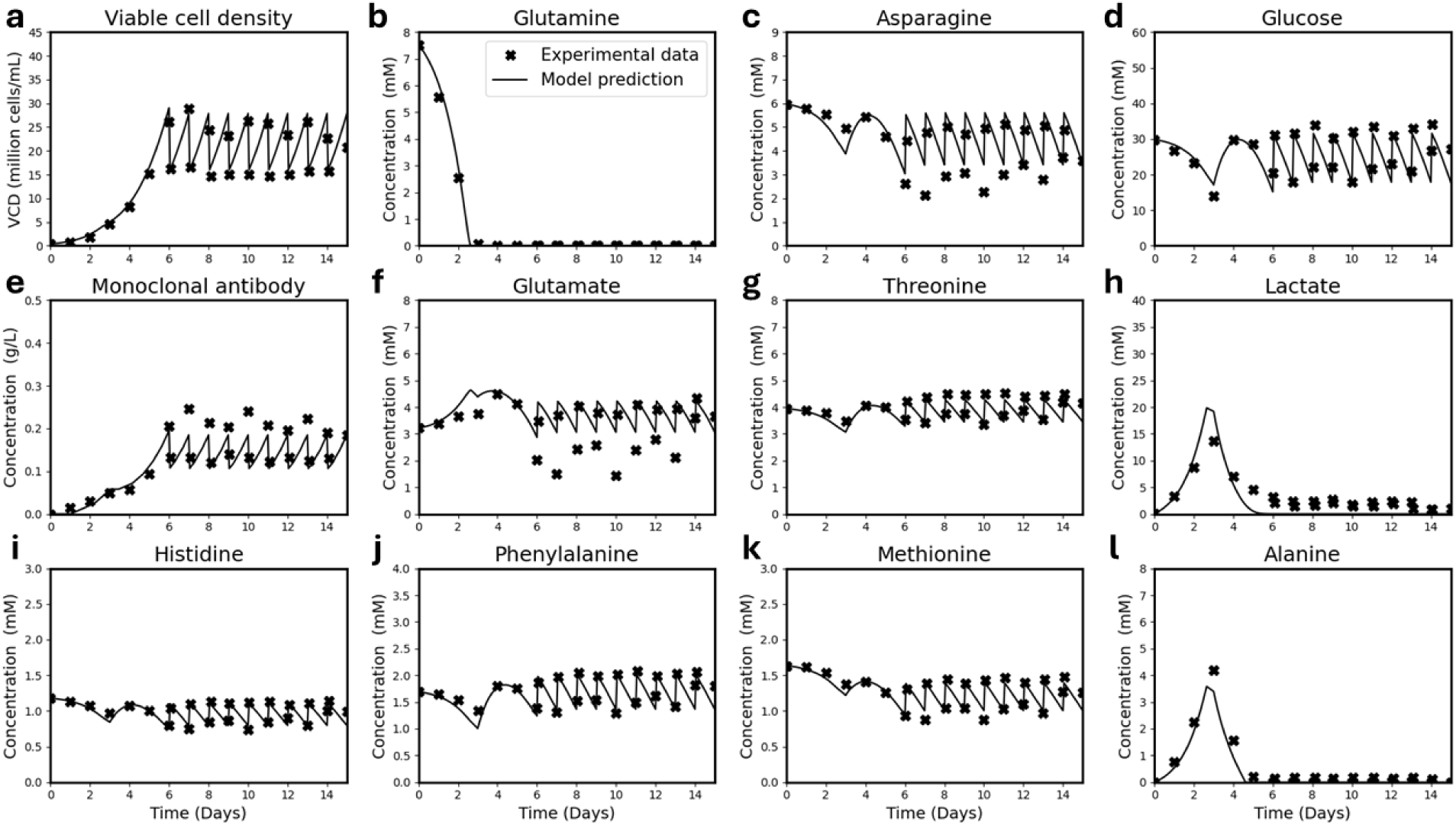
The model successfully predicts perfusion culture performance for concentrations of a) viable cell density, b) glutamine, c) asparagine, d) glucose, e) monoclonal antibody, f) glutamate, g) threonine, h) lactate, i) histidine, j) phenylalanine, k) methionine, and l) alanine.

### 4.4 Derisking cell culture media development

Another application of mathematical models is to guide development of cell culture media. Commercial media formulations are seldom reported in the literature. However, the composition of chemically defined AMBIC reference media has been published, leading to a more open-source approach to compare performance of bioprocesses. Data from fed-batch bioreactor cultures of the CHO VRC01 cell line in the AMBIC reference media has been published. This experiment involved seeding CHO VRC01 cells at 0.5 million cells/mL in the AMBIC basal media along with daily supplementation of AMBIC feed media. Measurements of cell density, glucose concentration, lactate concentration, glutamine concentration, glutamate concentration, and ammonia concentration were reported. (Cordova et al., 2023). We utilized this information along with the DMFA model to predict and validate the AMBIC media performance.

The model successfully predicts the peak viable cell density that can be achieved with the AMBIC media (Figure 7a). Figure 7 also compares the performance of the ActiPro and AMBIC media. The reason for the reduction of cell growth can be attributed to the depletion of asparagine (Figure 7c), valine (Figure 7j), and phenylalanine (Figure 7k). The predictions of glutamine, glutamate, ammonia, and mAb concentrations were also successful, highlighting that the model is capable of predicting culture performance under new media formulations prior to running experiments. Cell densities, glutamine, glutamate, ammonia, mAb, glucose, and lactate concentrations are commonly measured in cell culture experiments. Measurements of other amino acids are more difficult and expensive to measure and thus are not often reported. According to the literature data for the CHO VRC01 cell line, a peak viable cell density of 30 million cells was achieved using the ActiPro media with HyClone feeds, but the peak viable cell density was less than 10 million cells/mL with the AMBIC basal and feed media (Cordova et al., 2023). The successful predictions of viable cell density (Figure 7a) are enabled by the successful integration of different amino acids in the proposed model. These amino acids were present at higher quantities in the cultures grown with the ActiPro (with HyClone feed) media. Hence, we show that if a robust mathematical model is developed for a given cell line, the model can be used to provide potential explanations for poor bioprocess performance and predictions of culture performance under different media. Thus, highlighting that interpretability of mechanistic model predictions can aid with media development.

**Figure 7:**
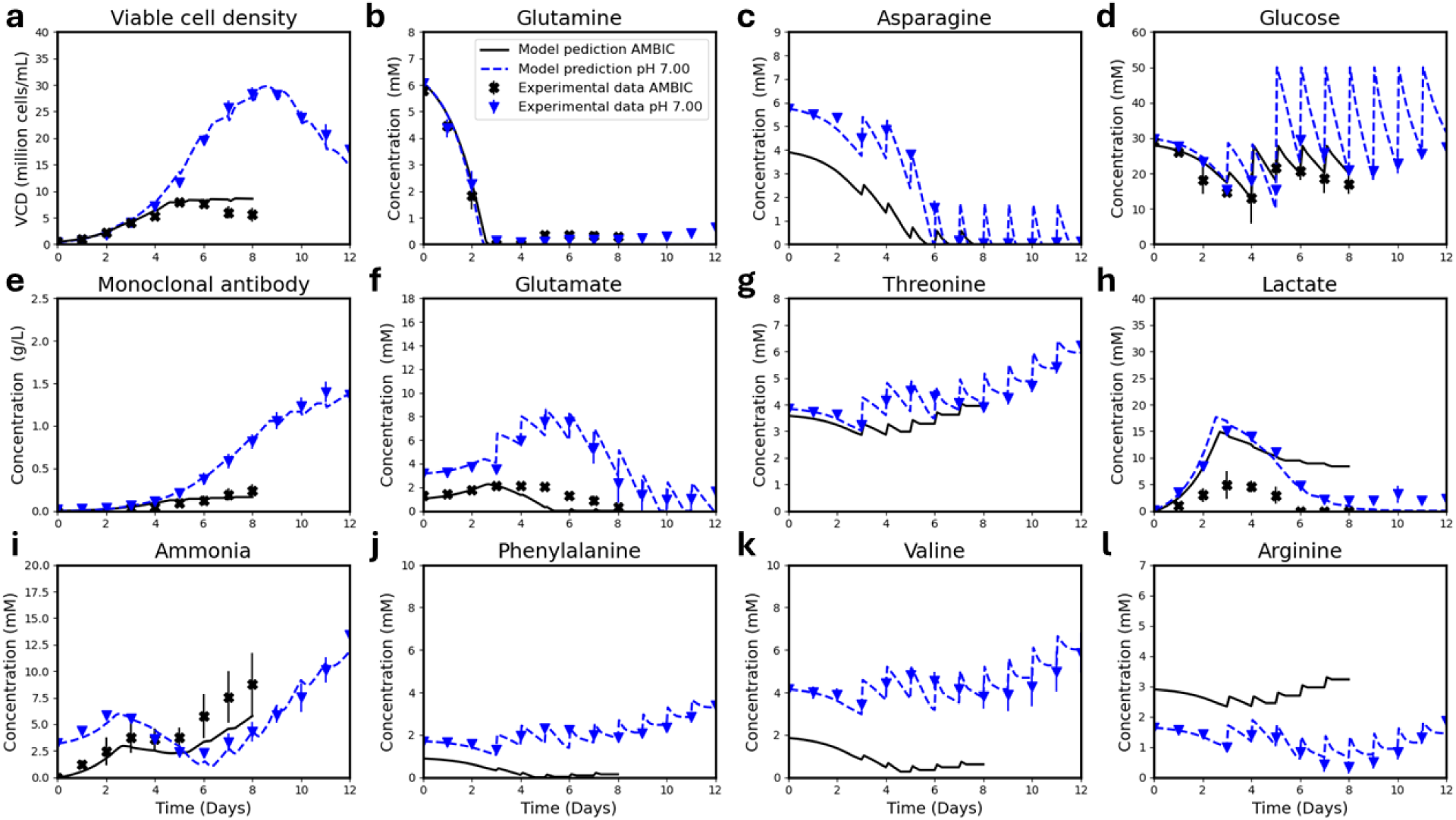
Predictions of CHO VRC01 fed-batch cultures while using the AMBIC basal and feed media along with experimental validation with literature experimental data. a) viable cell density, b) glutamine, c) asparagine, d) glucose, e) monoclonal antibody, f) glutamate, g) threonine, h) lactate, i) ammonia, j) phenylalanine, k) valine, and l) arginine.

## 5 Conclusions

The lack of detailed models of CHO cell metabolism that include the effect of bioreactor pH has been addressed in this work. Although dynamic metabolic flux analysis models exist in the literature (Nolan and Lee, 2011), they do not include the effect of bioreactor pH and also only model a subset of amino acids. Here, we developed a dynamic metabolic flux analysis model to predict concentrations of cell density, glucose, lactate, amino acids, ammonia, and mAb as a function of bioreactor pH, seeding cell density, and nutrient composition. This model was trained on experimental data from fed-batch experiments run at three different bioreactor pH conditions. The model was subsequently used to predict intensified fed-batch culture and perfusion culture performance. The model predictions were validated with experimental data thus demonstrating the model’s robust predictive capabilities. This work also shows the potential for model applications in industrially relevant process development scenarios, such as simulating intensified fed-batch and perfusion culture performance using only historical fed-batch data. Flowsheet models in the literature utilize mass balances to study the economic feasibility of fed-batch, intensified fed-batch, and perfusion bioreactors. Replacing these mass balances with mathematical models of cell metabolism, such as with the DMFA model developed in this study, can provide a more realistic flow sheet model to compare different operating conditions. The application of one model to predict performance of different bioreactor operation modes can help provide more realistic constraints to flow sheet models (Malinov et al., 2024). A survey of the application of digital twins in pharmaceutical manufacturing has highlighted that adaptive modeling methods with online streaming data need to be investigated (Chen et al., 2020). Due to ease of measurement cell culture processes typically involve online or at-line measurements of concentrations of glucose, lactate, mAb, ammonia, glutamine, glutamate, and cell density. Adaptive model calibration to this subset of online or at-line measurements can help realize the application of the DMFA model developed in this study as a digital twin capable of predicting concentrations of unmeasured amino acids by utilizing only the online or at-line measurements.

## Supporting information

Supplementary Section

## 6 Acknowledgements

This research was funded by US FOOD and DRUG ADMINISTRATION, grant number DHHS-FDA-R01FD006588, grant number U01FD007695A. This work was also supported with computational resources from the University of Delaware (Caviness cluster).

## 7 Nomenclature

G6P: Glucose-6-Phosphate
GLC: Glucose
ATP: Adenosine triphosphate
PYR: Pyruvate
NADH: Nicotinamide adenine dinucleotide (NAD)
GLU: Glutamic acid
ALA: Alanine
AKG: Alpha-ketoglutarate
LAC: Lactate
AcCoA: Acetyl-coenzyme A
CIT: Citrate
OXA: Oxaloacetate
SucCoA: Succinyl-coenzyme A
SUC: Succinate
FUM: Fumarate
TRP: Tryptophan
ASN: Asparagine
FADH_2_: Flavin adenine dinucleotide
ASP: Aspartic acid
PHE: Phenylalanine
TYR: Tyrosine
Acetoac: Acetoacetate
GLY: Glycine
SER: Serine
LEU: Leucine
HIS: Histidine
GLN: Glutamine
ARG: Arginine
.CYS: Cysteine
VAL: Valine
ILE: isoleucine
MET: Methionine
LYS: Lysine
THR: Threonine
ORN: Ornithine
CLN: Citrulline
AMBIC: Advanced mammalian biomanufacturing innovation center

## 8 Appendix A: Reaction network

**Table A1:** Metabolic reaction network used for metabolic flux analysis.

| Reaction number | Reaction | Rationale |
| --- | --- | --- |
| 1 | $\text{GLC} + \text{ATP} \rightarrow \text{G6P}$ | Intermediate steps in glycolysis were reduced to the listed reactions as additional reactions increase the complexity of the network with limited value. |
| 2 | $\text{G6P} \rightarrow 2 \text{PYR} + 2 \text{NADH} + 3 \text{ATP}$ | |
| 3 | $\text{PYR} + \text{NADH} \rightleftharpoons \text{LAC}$ | |
| 4 | $\text{PYR} + \text{GLU} \rightleftharpoons \text{ALA} + \text{AKG}$ | |
| 5 | $\text{PYR} \rightarrow \text{AcCoA} + \text{CO}_2 + \text{NADH}$ | The original reaction network consisted of the reaction from Fumarate to Malate. This reaction was removed as it was a linear reaction, and Fumarate can replace malate in subsequent reactions for the purpose of modeling. |
| 6 | $\text{AcCoA} + \text{OXA} \rightarrow \text{CIT}$ | |
| 7 | $\text{CIT} \rightarrow \text{CO}_2 + \text{NADH} + \text{AKG}$ | |
| 8 | $\text{AKG} \rightarrow \text{NADH} + \text{CO}_2 + \text{SucCoA}$ | |
| 9 | $\text{SucCoA} \rightleftharpoons \text{ATP} + \text{Suc}$ | |
| 10 | $\text{Suc} \rightleftharpoons \text{Fum} + \text{FADH}_2$ | |
| 11 | $\text{Fum} \rightarrow \text{PYR} + \text{CO}_2$ | |
| 12 | $\text{Fum} \rightleftharpoons \text{OXA} + \text{NADH}$ | |
| 13 | $\text{TRP} \rightarrow 2 \text{AcCoA} + 4 \text{CO}_2 + 2 \text{NADH} + 1 \text{ALA}$ | Minimal changes were made to amino acid metabolism reactions. (Zamorano et al., 2010) consisted of the following reactions<br>$\alpha\text{Ka} \rightarrow \text{AcetoAcCoA} + 2\text{NADH} + 2\text{CO}_2$<br>$\text{AcetoAcCoA} \rightarrow 2\text{AcCoA}$<br>These reactions were collapsed to yield:<br>$\alpha\text{Ka} \rightarrow 2\text{AcCoA} + 2\text{NADH} + 2\text{CO}_2$<br>Thus, $\alpha\text{Ka}$ was replaced with these products.<br>The following reaction was also removed:<br>$\text{PropCoA} + \text{ATP} \rightarrow \text{SucCoA}$<br>PropCoA was removed and replaced with ATP and SucCoA with the |
| 14 | $\text{ASN} \rightleftharpoons \text{NH}_3 + \text{ASP}$ | |
| 15 | $\text{ASP} + \text{AKG} \rightleftharpoons \text{OXA} + \text{GLU}$ | |
| 16 | $\text{PHE} + \text{NADH} \rightarrow \text{TYR}$ | |
| 17 | $\text{TYR} + \text{AKG} \rightarrow \text{GLU} + \text{CO}_2 + \text{Acetoac} + \text{FUM}$ | |
| 18 | $\text{SER} + \text{NH}_3 + \text{NADH} + \text{CO}_2 \rightleftharpoons 2 \text{GLY}$ | |
| 19 | $\text{LEU} + \text{AKG} + \text{ATP} \rightarrow \text{GLU} + \text{Acetoac} + \text{NADH} + \text{FADH}_2 + \text{AcCoA}$ | |
| 20 | $\text{THR} \rightarrow \text{GLY} + \text{NADH} + \text{AcCoA}$ | |
| 21 | $\text{HIS} \rightarrow \text{GLU} + \text{NH}_3$ | |
| 22 | $\text{GLN} \rightleftharpoons \text{GLU} + \text{NH}_3$ | |
| 23 | $\text{GLU} \rightleftharpoons \text{AKG} + \text{NH}_3 + \text{NADH}$ | |
| 24 | $\text{CYS} \rightarrow \text{PYR} + \text{NH}_3$ | |
| 25 | $\text{VAL} + \text{ATP} + \text{AKG} \rightarrow \text{GLU} + 2\text{CO}_2 + 3\text{NADH} + \text{FADH}_2 + \text{SucCoA}$ | |
| 26 | $\text{ILE} + \text{ATP} + \text{AKG} \rightarrow \text{GLU} + \text{AcCoA} + 2 \text{NADH} + \text{FADH}_2 + \text{SucCoA}$ | |
| 27 | $\text{THR} \rightleftharpoons \text{NADH} + \text{NH}_3 + \text{Suc}$ | appropriate stoichiometry in subsequent reactions.<br>The following reaction was also removed:<br>$\text{Glu}\gamma\text{SA} \rightarrow \text{GLU}$<br>$\text{Glu}\gamma\text{SA}$ was replaced with $\text{GLU}$ in subsequent reactions.<br>The following reaction was also removed:<br>$\text{ArgSucc} \rightarrow \text{Arg} + \text{Fum}$<br>$\text{ArgSucc}$ was replaced with $\text{Arg}$ and $\text{Fum}$ with the appropriate stoichiometry in subsequent reactions. |
| 28 | $\text{MET} + \text{SER} + 2\text{ATP} \rightarrow \text{CYS} + \text{NADH} + \text{SucCoA} + \text{NH}_3$ | |
| 29 | $\text{Acetoac} + \text{SucCoA} \rightarrow 2 \text{AcCoA} + \text{Suc}$ | |
| 30 | $2 \text{AKG} + \text{LYS} \rightarrow 2 \text{GLU} + 2 \text{AcCoA} + 3 \text{NADH} + 1 \text{FADH}_2$ | |
| 31 | $1 \text{NADH} + 0.5 \text{O}_2 \rightarrow 2.5 \text{ATP}$ | |
| 32 | $1 \text{FADH}_2 + 0.5 \text{O}_2 \rightarrow 1.5 \text{ATP}$ | |
| 33 | $\text{ARG} \rightarrow \text{ORN} + \text{Urea}$ | |
| 34 | $\text{ORN} + \text{NH}_3 + \text{CO}_2 + 2\text{ATP} \rightarrow \text{CLN}$ | |
| 35 | $\text{ASP} + \text{CLN} + \text{ATP} \rightarrow \text{FUM} + \text{ARG}$ | |
| 36 | $\text{ORN} + \text{AKG} \rightleftharpoons 2 \text{GLU} + \text{NADH}$ | |
| 37 | $\text{G6P} + 2 \text{GLU} \rightarrow 2 \text{SER} + 2 \text{AKG} + \text{NADH}$ | |
| 38 | $\text{SER} \rightarrow \text{PYR} + \text{NH}_3$ | |
| 39 | $0.0077 \text{ALA} + 0.0063 \text{ARG} + 0.0046 \text{ASN} + 0.0119 \text{ASP} + 0.0020 \text{CYS} + 0.0167 \text{GLN} + 0.0126 \text{GLY} + 0.0020 \text{HIS} + 0.0038 \text{ILE} + 0.0082 \text{LEU} + 0.0072 \text{LYS} + 0.0018 \text{MET} + 0.0034 \text{PHE} + 0.0071 \text{SER} + 0.0052 \text{THR} + 0.001 \text{TRP} + 0.0028 \text{TYR} + 0.0050 \text{VAL} + 0.5444 \text{ATP} + 0.0025 \text{NH}_3 + 0.1178 \text{NADH} + 0.0843 \text{AcCoA} + 0.0123 \text{G6P} \rightarrow 0.0132 \text{CO}_2 + 0.0041 \text{GLU} + 0.0063 \text{FUM} + 1.0 \text{Biomass}$ | (Zamorano et al., 2010)) consisted of reactions involving synthesis of DNA and RNA from nucleotides, biomass protein from amino acids, and membrane lipids from amino acids and lipid precursors.<br>The ratios of membrane lipids, DNA, RNA, and protein required to produce a Carbon mol of Biomass can vary drastically (Szélieová et al., 2020).<br>(Zamorano et al., 2010)consisted of the following reaction.<br>Reactions with very small or very large stoichiometric coefficients can lead to ill conditions matrices. Hence the reactions producing RNA, DNA, Protein, membrane lipids, and the overall biomass reaction from various metabolites were lumped to |
|  |  | yield one reaction. The calculations are in the section below. |
| 40 | $\begin{aligned} &0.0096 \text{ ALA} + 0.0083 \text{ ARG} + 0.008 \text{ ASN} \\ &+ 0.0087 \text{ ASP} + 0.0056 \text{ CYS} + 0.0105 \\ &\text{GLU} + 0.0096 \text{ GLN} + 0.0142 \text{ GLY} + \\ &0.0037 \text{ HIS} + 0.0053 \text{ ILE} + 0.0148 \text{ LEU} + \\ &0.0127 \text{ LYS} + 0.0022 \text{ MET} + 0.0068 \text{ PHE} \\ &+ 0.0232 \text{ SER} + 0.0152 \text{ THR} + 0.004 \\ &\text{TRP} + 0.0087 \text{ TYR} + 0.0182 \text{ VAL} + \\ &0.816 \text{ ATP} \rightarrow \text{IgG} \end{aligned}$ | The amino acid sequence of the mAb was used to determine the amino acid fraction required to produce 1 Carbon mol (normalized by number of carbons) of IgG.<br>The molecular weight of the aglycosylated mAb is 145755.29 Da and the molecular weight per carbon mol is 22.54. Hence, while performing flux balance analysis, the<br>$q_{mAb} = q_{IgG} \times \frac{145755.29}{22.54}$ |
| 41 | $\text{Urea} \rightleftharpoons \text{Urea}_{\text{ext}}$ | These are exchange reactions that dictate secretion and intake of metabolites across the cell membrane. |
| 42 | $\text{ATP} \rightarrow \text{ATP}_{\text{ext}}$ | |
| 43 | $\text{CYS}_{\text{ext}} \rightarrow \text{CYS}$ | |
| 44 | $\text{O}_{2,\text{ext}} \rightarrow \text{O}_2$ | |
| 45 | $\text{CO}_{2,\text{ext}} \rightleftharpoons \text{CO}_2$ | |
| 46 | $\text{LAC}_{\text{ext}} \rightleftharpoons \text{LAC}$ | |
| 47 | $\text{GLC}_{\text{ext}} \rightarrow \text{GLC}$ | |
| 48 | $\text{ALA}_{\text{ext}} \rightleftharpoons \text{ALA}$ | |
| 49 | $\text{TRP}_{\text{ext}} \rightarrow \text{TRP}$ | |
| 50 | $\text{NH}_{3,\text{ext}} \rightleftharpoons \text{NH}_3$ | |
| 51 | $\text{ASN}_{\text{ext}} \rightarrow \text{ASN}$ | |
| 52 | $\text{ASP}_{\text{ext}} \rightleftharpoons \text{ASP}$ | |
| 53 | $\text{PHE}_{\text{ext}} \rightarrow \text{PHE}$ | |
| 54 | $\text{TYR}_{\text{ext}} \rightarrow \text{TYR}$ | |
| 55 | $\text{SER}_{\text{ext}} \rightleftharpoons \text{SER}$ | |
| 56 | $\text{GLY}_{\text{ext}} \rightleftharpoons \text{GLY}$ | |
| 57 | $\text{MET}_{\text{ext}} \rightarrow \text{MET}$ | |
| 58 | $\text{THR}_{\text{ext}} \rightarrow \text{THR}$ | |
| 59 | $\text{ILE}_{\text{ext}} \rightarrow \text{ILE}$ | |
| 60 | $\text{VAL}_{\text{ext}} \rightarrow \text{VAL}$ | |
| 61 | $\text{GLU}_{\text{ext}} \rightleftharpoons \text{GLU}$ | |
| 62 | $\text{ARG}_{\text{ext}} \rightarrow \text{ARG}$ | |
| 63 | $\text{GLN}_{\text{ext}} \rightleftharpoons \text{GLN}$ | |
| 64 | $\text{HIS}_{\text{ext}} \rightarrow \text{HIS}$ | |
| 65 | $\text{LEU}_{\text{ext}} \rightarrow \text{LEU}$ | |
| 66 | $\text{LYS}_{\text{ext}} \rightarrow \text{LYS}$ | |
| 67 | $\text{Biomass}_{\text{ext}} \rightarrow \text{Biomass}$ | |
| 68 | $\text{IgG}_{\text{ext}} \rightarrow \text{IgG}$ | |

## 9 Appendix B: Kinetic parameter description and values

**Table B1:** Common kinetic parameter nomenclature and values.

| Parameter | Name | Value | Units |
| --- | --- | --- | --- |
| **Growth and death** |  |  |  |
| $K_{S,GLC}$ | Glucose Monod constant | 4.007 | mM |
| $K_{S,GLN}$ | Glutamine Monod constant | 0.239 | mM |
| $K_{S,GLU}$ | Glutamate Monod constant | 0.001 | mM |
| $K_{S,ASN}$ | Asparagine Monod constant | 1.000 | mM |
| $K_{S,ASP}$ | Aspartate Monod constant | 0.109 | mM |
| $K_{S,MET}$ | Methionine Monod constant | 0.001 | mM |
| $kbiphasic_{GLN}$ | Maximum specific growth rate on glutamine | 0.344 | day<sup>-1</sup> |
| $kbiphasic_{GLU}$ | Maximum specific growth rate on glutamate | 0.656 | day<sup>-1</sup> |
| $kbiphasic_{ASN}$ | Maximum specific growth rate on asparagine | 0.858 | day<sup>-1</sup> |
| $kbiphasic_{ASP}$ | Maximum specific growth rate on aspartate | 0.142 | day<sup>-1</sup> |
| $OSMO\_A$ | Osmolarity at half the maximum specific growth rate | 429.598 | mOsm kg<sup>-1</sup> |
| $OSMO\_B$ | Rate at which the specific growth rate decreases with increasing osmolality | 0.072 | kg mOsm<sup>-1</sup> |
| $\mu_{death,1}$ | Constant specific death rate | 0.01 | day<sup>-1</sup> |
| $\mu_{death,2}$ | Maximum specific death rate due to accumulation of unmeasurable inhibitory metabolites | 0.338 | day<sup>-1</sup> |
| $P_{1,death,IVCD}$ | IVCD at half of the maximum specific death rate | 120 | (10<sup>6</sup> cells) day mL<sup>-1</sup> |
| $P_{2,death}$ | Rate at which the specific death rate increases with increasing IVCD | 0.100 | mL (10<sup>6</sup> cells)<sup>-1</sup> day<sup>-1</sup> |
| **Substrates** |  |  |  |
| $m_{GLC}$ | Glucose maintenance coefficient | 918.109 | nmol (10<sup>6</sup> cells)<sup>-1</sup> day<sup>-1</sup> |
| $Y_{\frac{X}{LAC}}$ | Yield of cells on lactate | 9.881 | nmol cells nmol<sup>-1</sup> |
| $K_{S,LAC}$ | Lactate Monod constant | 2.731 | mM |
| $Y_{\frac{X}{ALA}}$ | Yield of cells on alanine | 36.695 | nmol cells nmol<sup>-1</sup> |
| $Y_{\frac{X}{SER}}^A$ | Yield of cells on serine during early culture | 8.684 | nmol cells nmol<sup>-1</sup> |
| $Y_{\frac{X}{SER}}^B$ | Yield of cells on serine during late culture | 17.621 | nmol cells nmol<sup>-1</sup> |
| $Y_{\frac{X}{GLY}}$ | Yield of cells on glycine | 300.543 | nmol cells nmol<sup>-1</sup> |
| $Y_{\frac{X}{GLN}}$ | Yield of cells on glutamine | 1.327 | nmol cells nmol<sup>-1</sup> |
| $m_{GLU}$ | Glutamine maintenance coefficient | 156.979 | nmol (10<sup>6</sup> cells)<sup>-1</sup> day<sup>-1</sup> |
| $Y_{\frac{X}{ASN}}$ | Yield of cells from asparagine | 22.209 | nmol cells nmol<sup>-1</sup> |
| $m_{ASN}$ | Asparagine maintenance coefficient | 115.079 | nmol (10<sup>6</sup> cells)<sup>-1</sup> day<sup>-1</sup> |
| $m_{ASP}$ | Aspartate maintenance coefficient | 105.692 | nmol (10<sup>6</sup> cells)<sup>-1</sup> day<sup>-1</sup> |
| $Y_{\frac{X}{MET}}$ | Yield of cells on methionine | 90.708 | nmol cells nmol<sup>-1</sup> |
| $Y_{\frac{X}{TYR}}$ | Yield of cells on tyrosine | 73.629 | nmol cells nmol<sup>-1</sup> |
| $Y_{\frac{X}{TRP}}$ | Yield of cells on tryptophan | 151.565 | nmol cells nmol<sup>-1</sup> |
| $Y_{\frac{X}{PHE}}$ | Yield of cells on phenylalanine | 54.739 | nmol cells nmol<sup>-1</sup> |
| $Y_{\frac{X}{VAL}}$ | Yield of cells on valine | 30.000 | nmol cells nmol<sup>-1</sup> |
| $Y_{\frac{X}{LYS}}$ | Yield of cells on lysine | 40.051 | nmol cells nmol<sup>-1</sup> |
| $Y_{\frac{X}{LEU}}$ | Yield of cells on leucine | 25.467 | nmol cells nmol<sup>-1</sup> |
| $Y_{\frac{X}{ILE}}$ | Yield of cells on isoleucine | 39.378 | nmol cells nmol<sup>-1</sup> |
| $Y_{\frac{X}{ARG}}$ | Yield of cells on arginine | 53.992 | nmol cells nmol<sup>-1</sup> |
| $Y_{\frac{X}{HIS}}$ | Yield of cells on histidine | 111.776 | nmol cells nmol<sup>-1</sup> |
| $Y_{\frac{X}{THR}}$ | Yield of cells on threonine | 42.504 | nmol cells nmol<sup>-1</sup> |
| $R_{AMM,1}$ | Ammonia consumed by CHO cells for glutamine/asparagine synthesis | 0.068 | mol (10<sup>6</sup> cells)<sup>-1</sup> |
| **Substrate metabolism interconversion** |  |  |  |
| $Y_{\frac{ALA}{GLC}}$ | Yield of alanine from glucose | 0.363 | nmol nmol<sup>-1</sup> |
| $Y_{\frac{GLY}{SER}}$ | Yield of glycine from serine | 0.728 | nmol nmol<sup>-1</sup> |
| $Y_{\frac{ASP}{GLU}}$ | Yield of aspartate from glutamine | 0.100 | nmol nmol<sup>-1</sup> |
| $Y_{\frac{GLU}{GLN}}$ | Yield of glutamate from glutamine | 0.200 | nmol nmol<sup>-1</sup> |
| $Y_{\frac{AMM}{GLN}}$ | Yield of ammonia from glutamine | 0.650 | nmol nmol<sup>-1</sup> |

**Table B2:** Parameters as a quadratic function of bioreactor pH.

| Parameter | Name | A | B | C | Units |
| --- | --- | --- | --- | --- | --- |
| $Y_{\frac{X}{GLC}}$ | Yield of cells on glucose | -13.435 | 180.598 | -600.754 | nmol cells nmol<sup>-1</sup> |
| $\mu_{max}$ | Maximum specific growth rate | -5.222 | 73.572 | -257.851 | day<sup>-1</sup> |
| $Y_{\frac{LAC}{GLC}}$ | Yield of lactate from glucose | -7.595 | 108.228 | -383.444 | nmol nmol<sup>-1</sup> |
| $q_{mAb}$ | mAb production rate | 50.909 | 194.225 | -3381.181 | nmol (10<sup>6</sup> cells)<sup>-1</sup> day<sup>-1</sup> |
| *AA scaling ratio* | Scaling ratio for amino acid demands | -1.44 | 19.8 | -67.04 | Unitless |
| $R_{AMM,2}$ | Lumped ammonia secretion from net amino acid catabolism | 212.237 | -<br>3166.862 | 11935.112 | nmol (10<sup>6</sup> cells)<sup>-1</sup> day<sup>-1</sup> |

## 10 Appendix C: Kinetic equations and mass balances

**Table C1:**
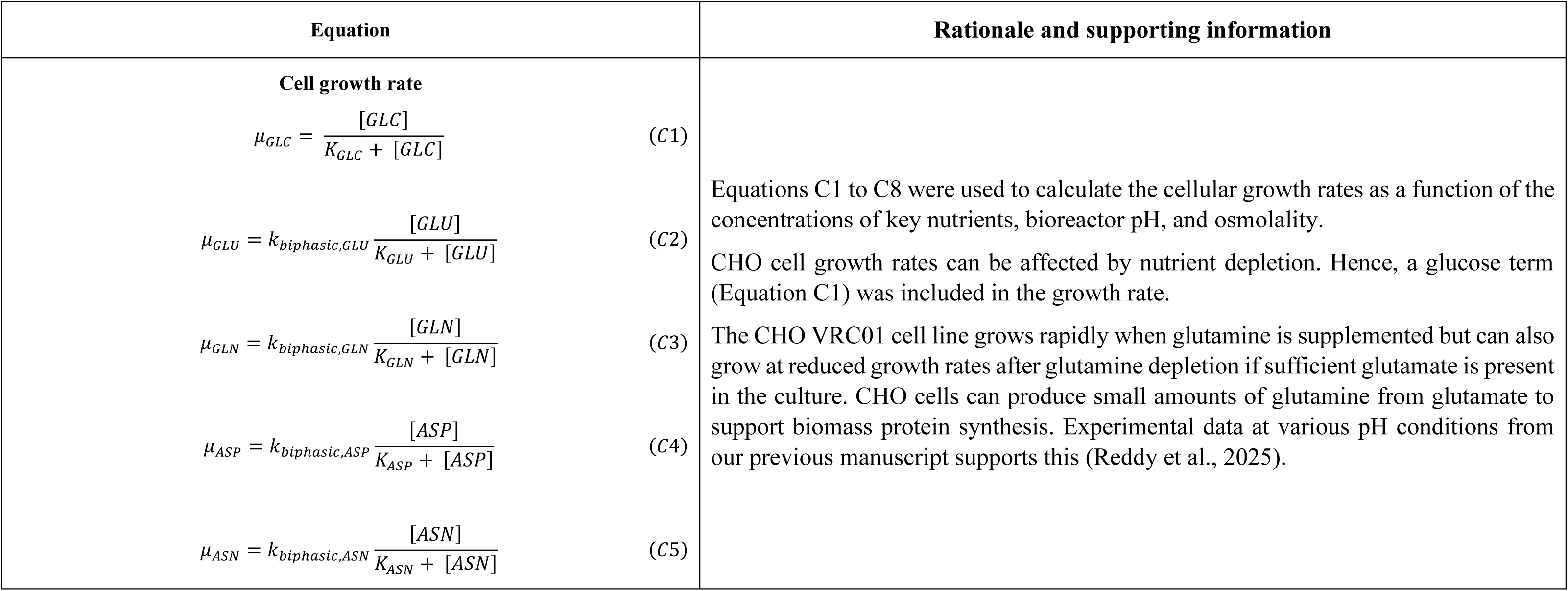
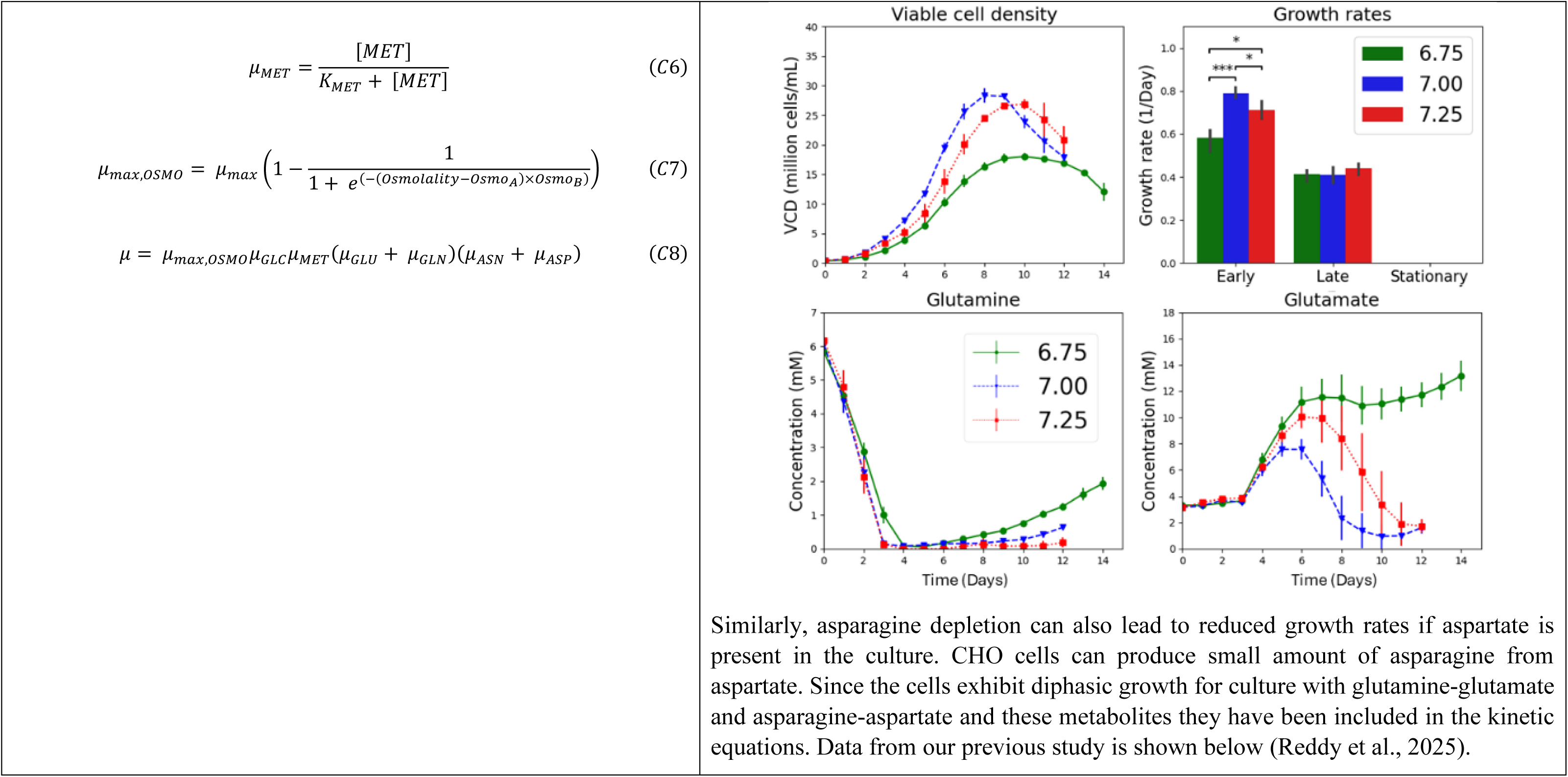
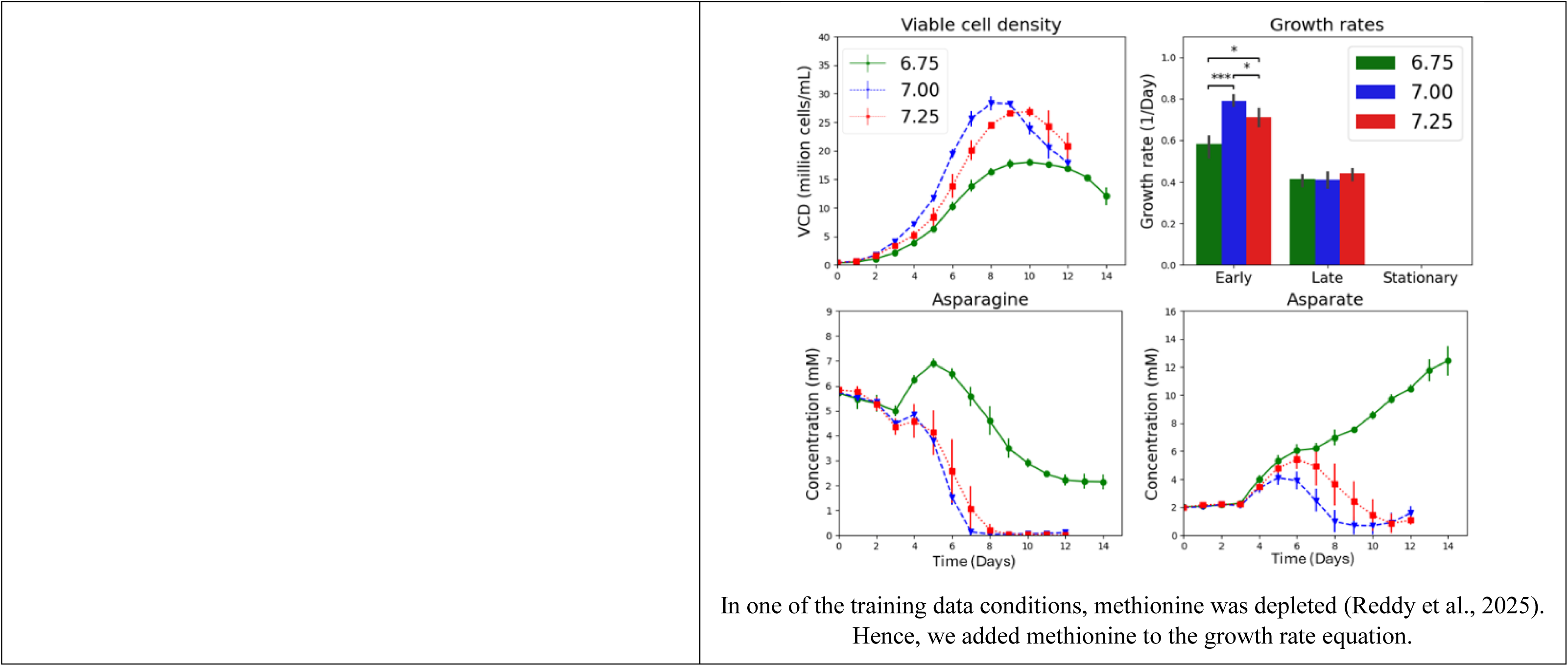
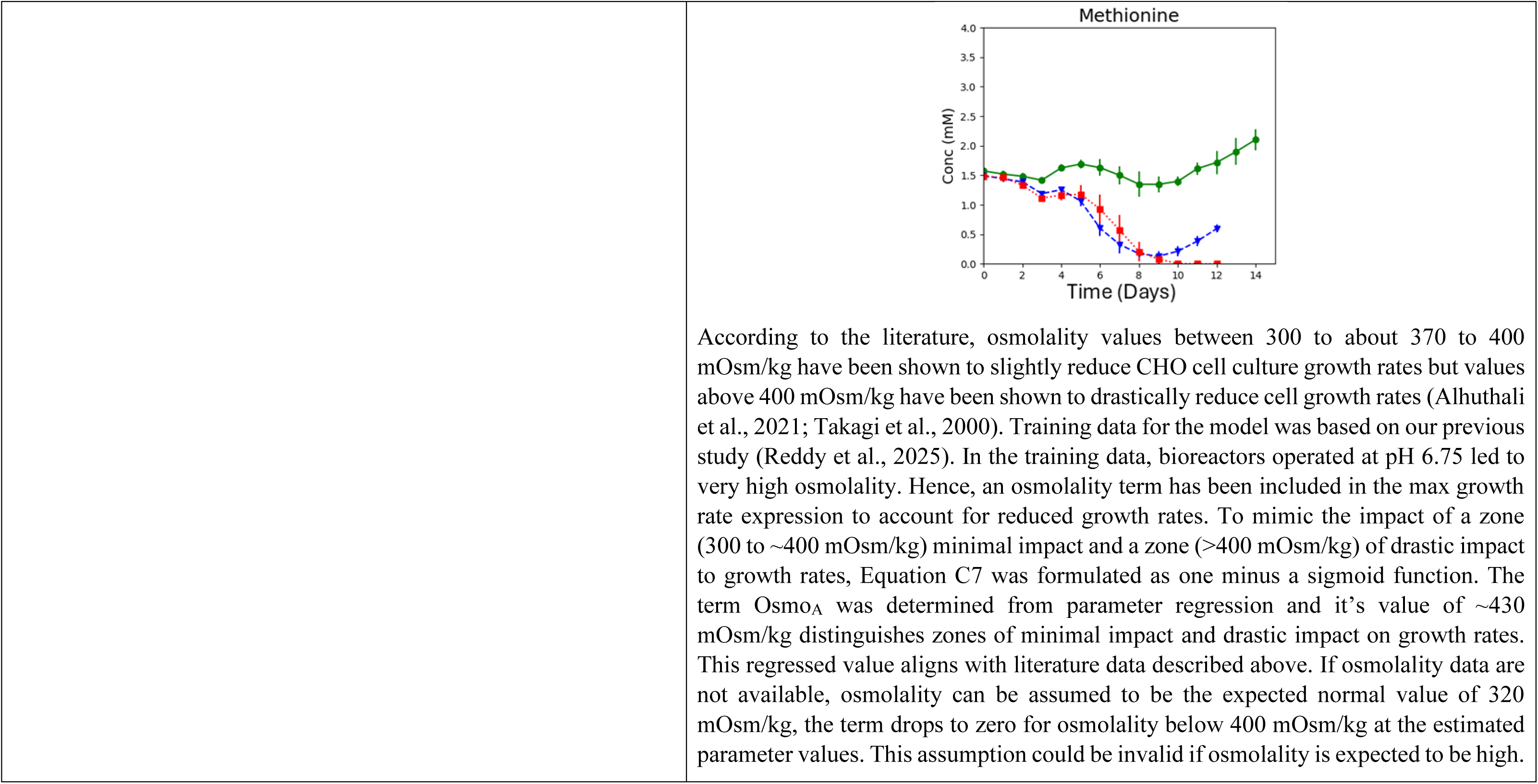
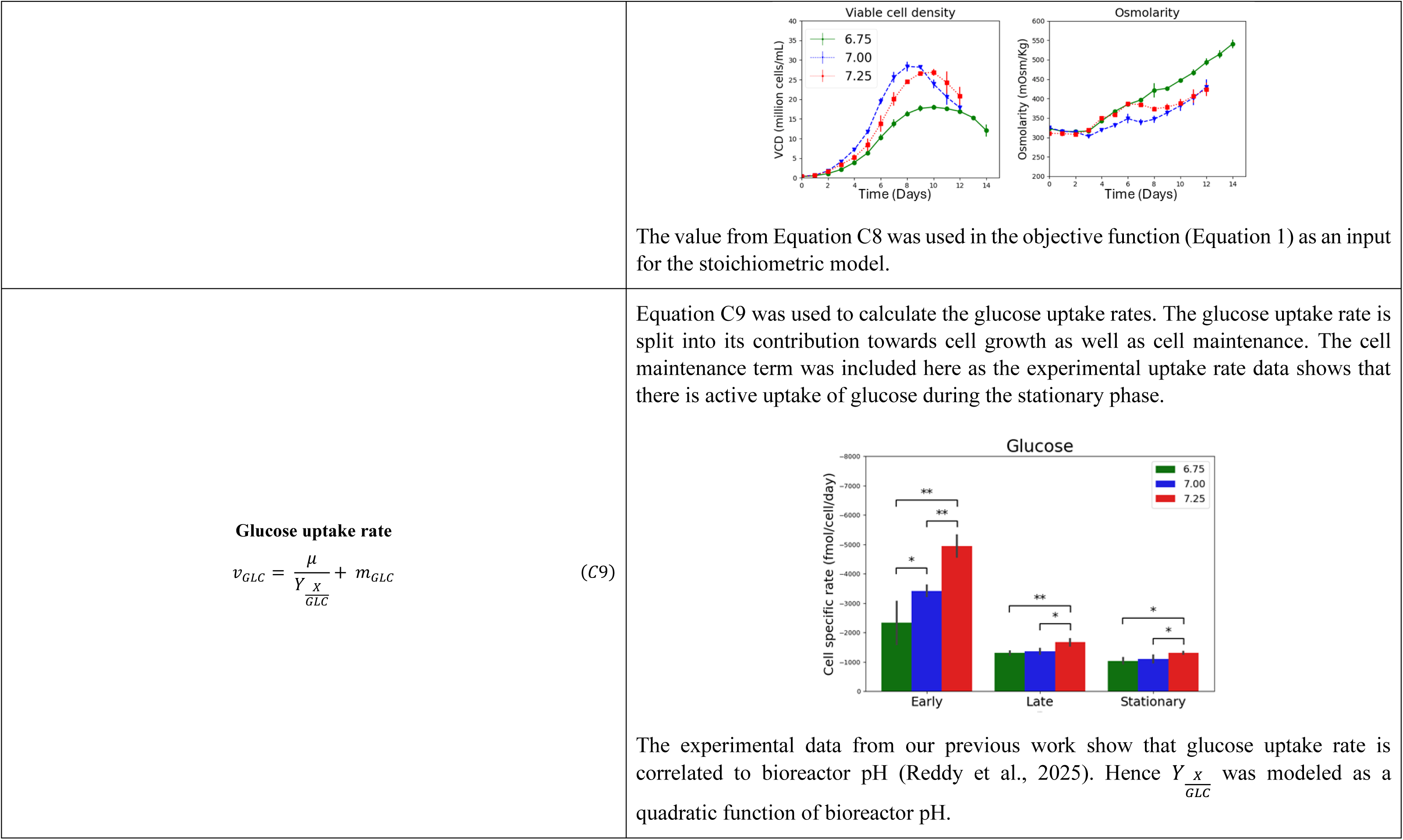
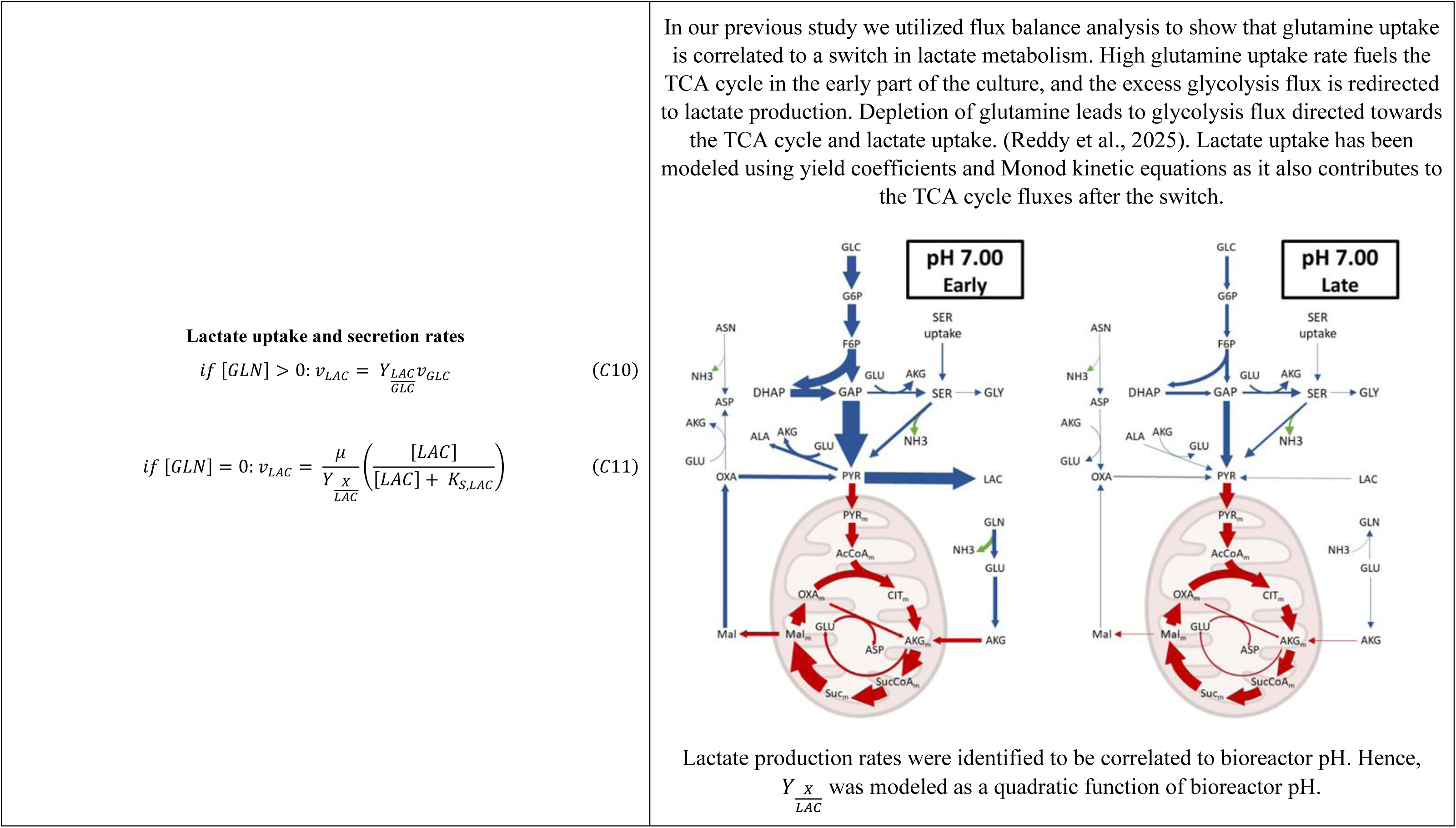
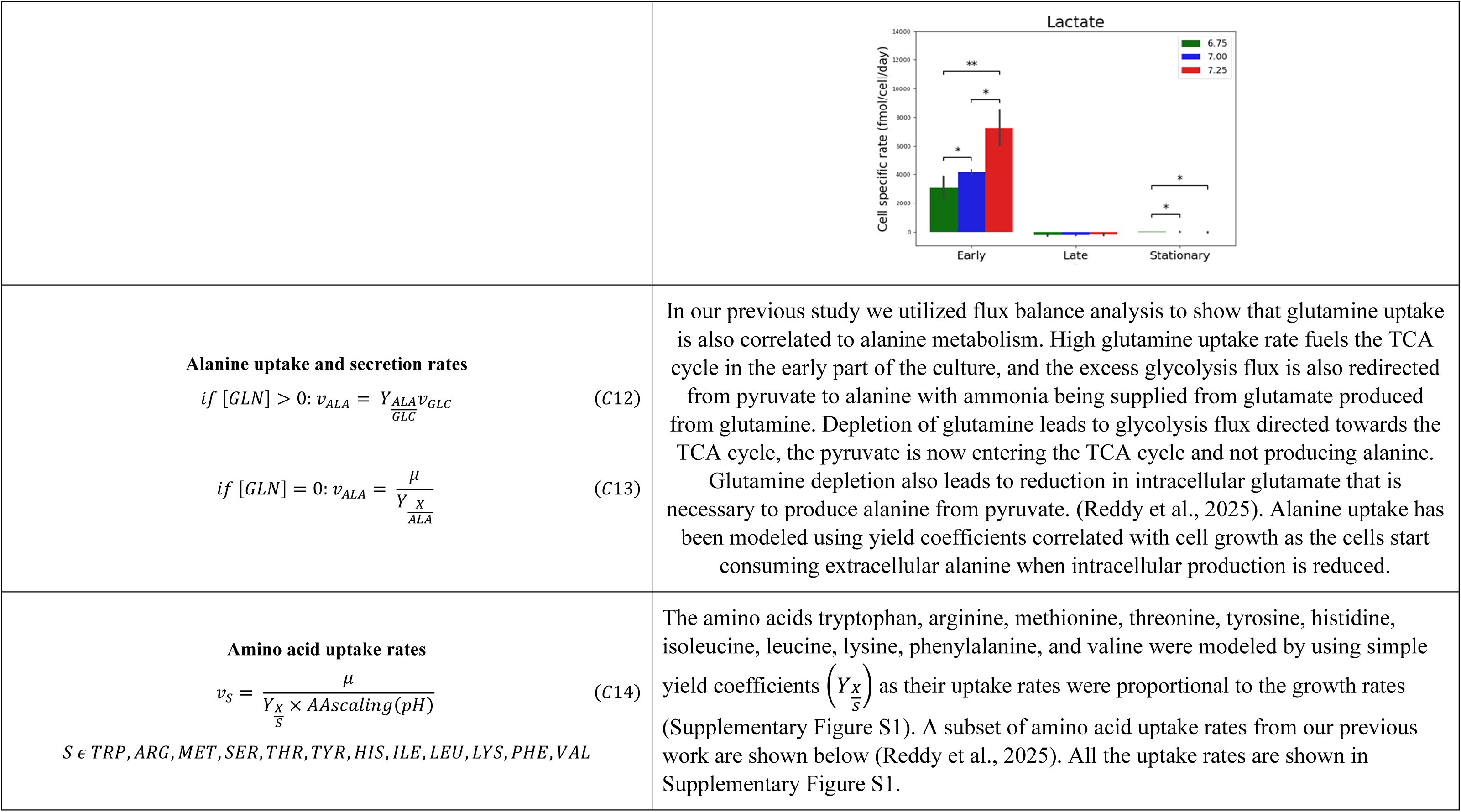
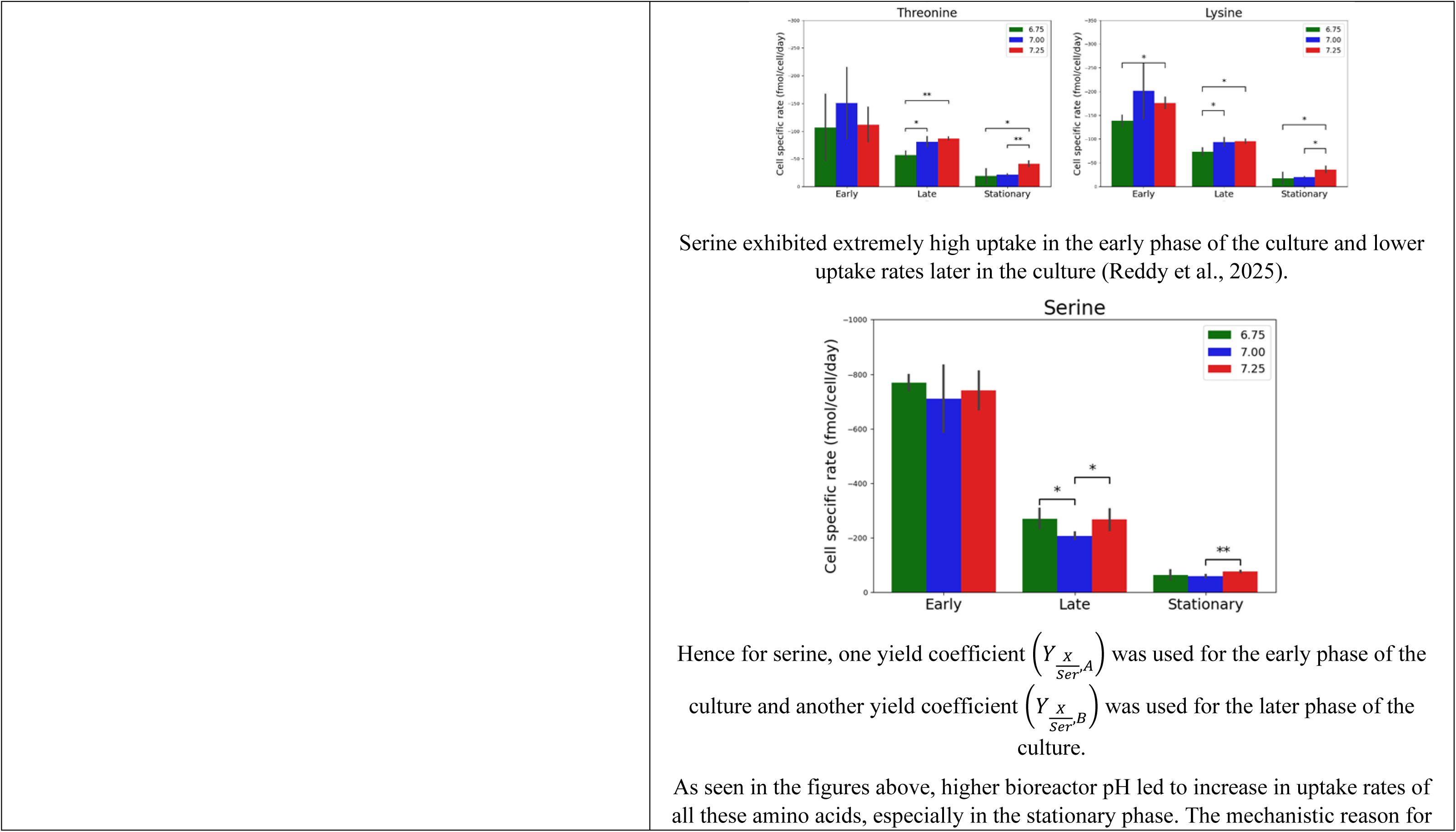
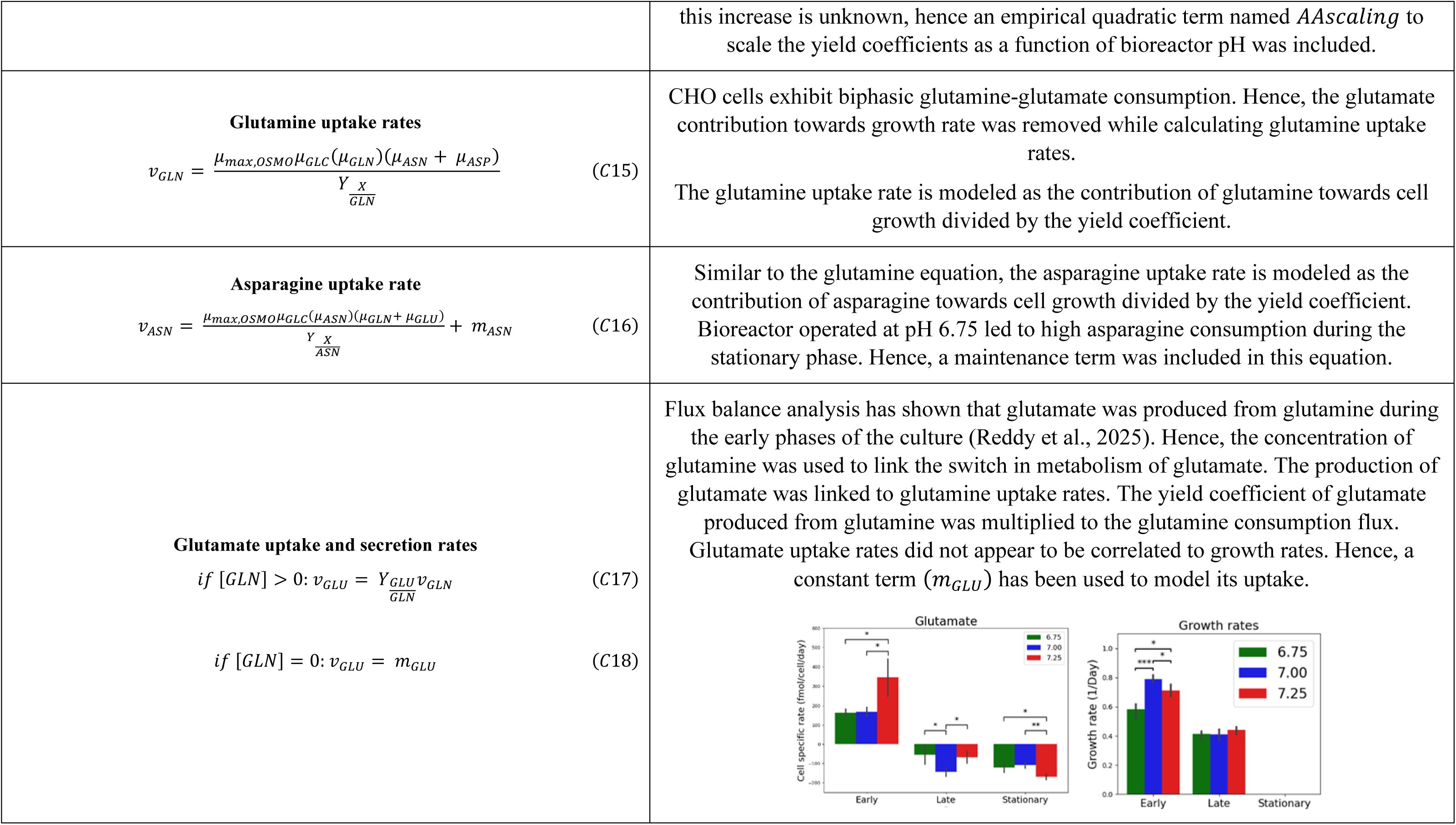
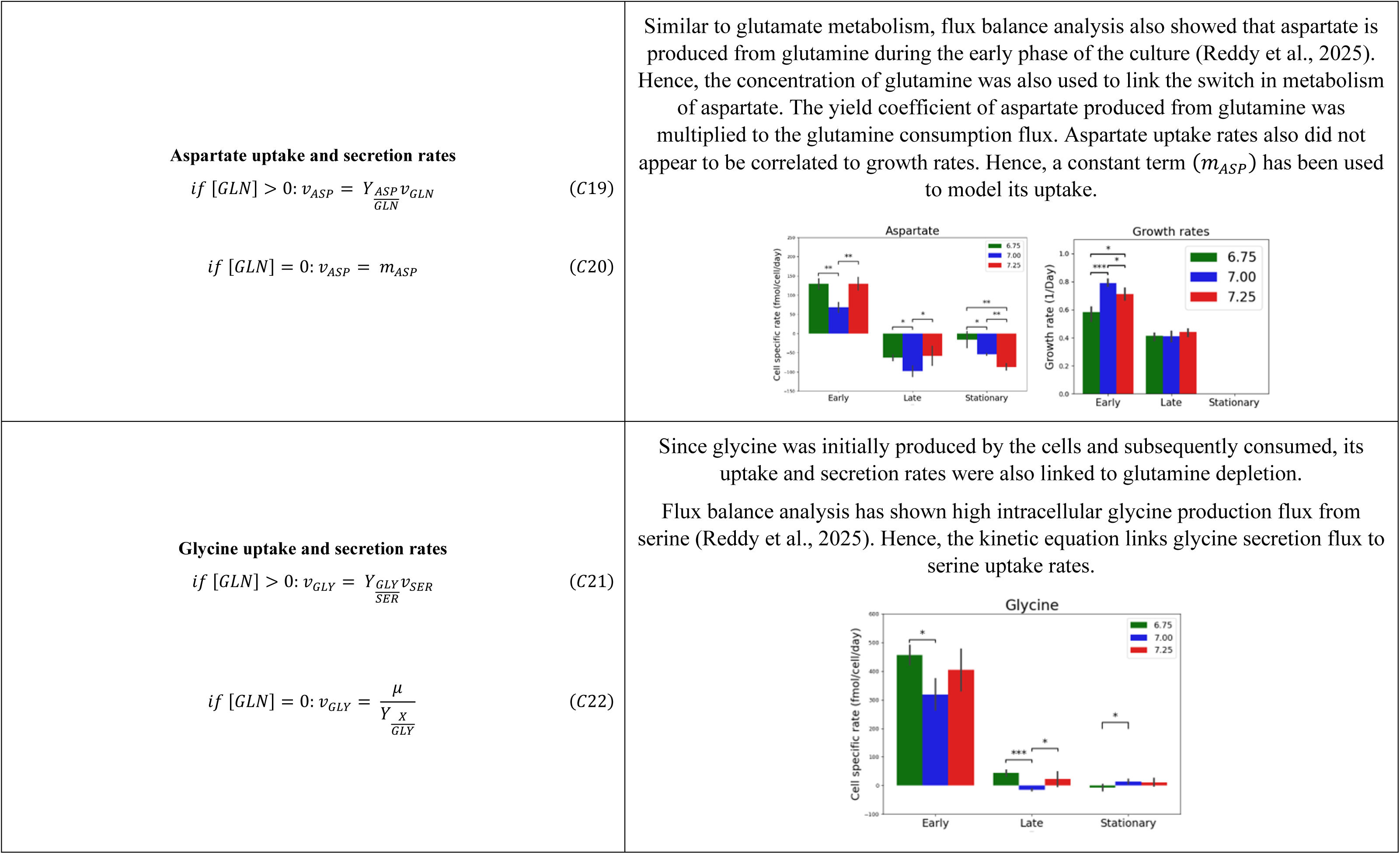
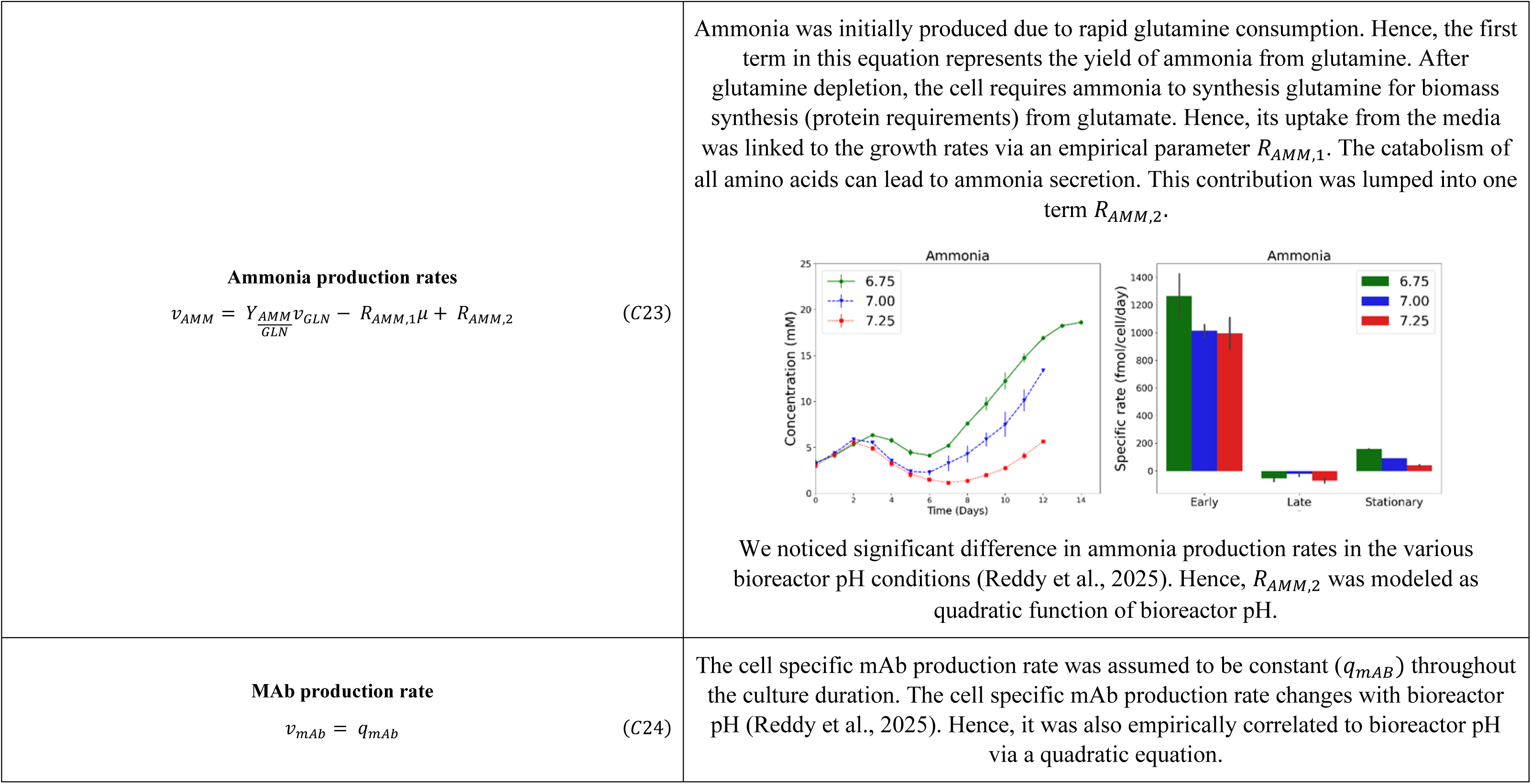
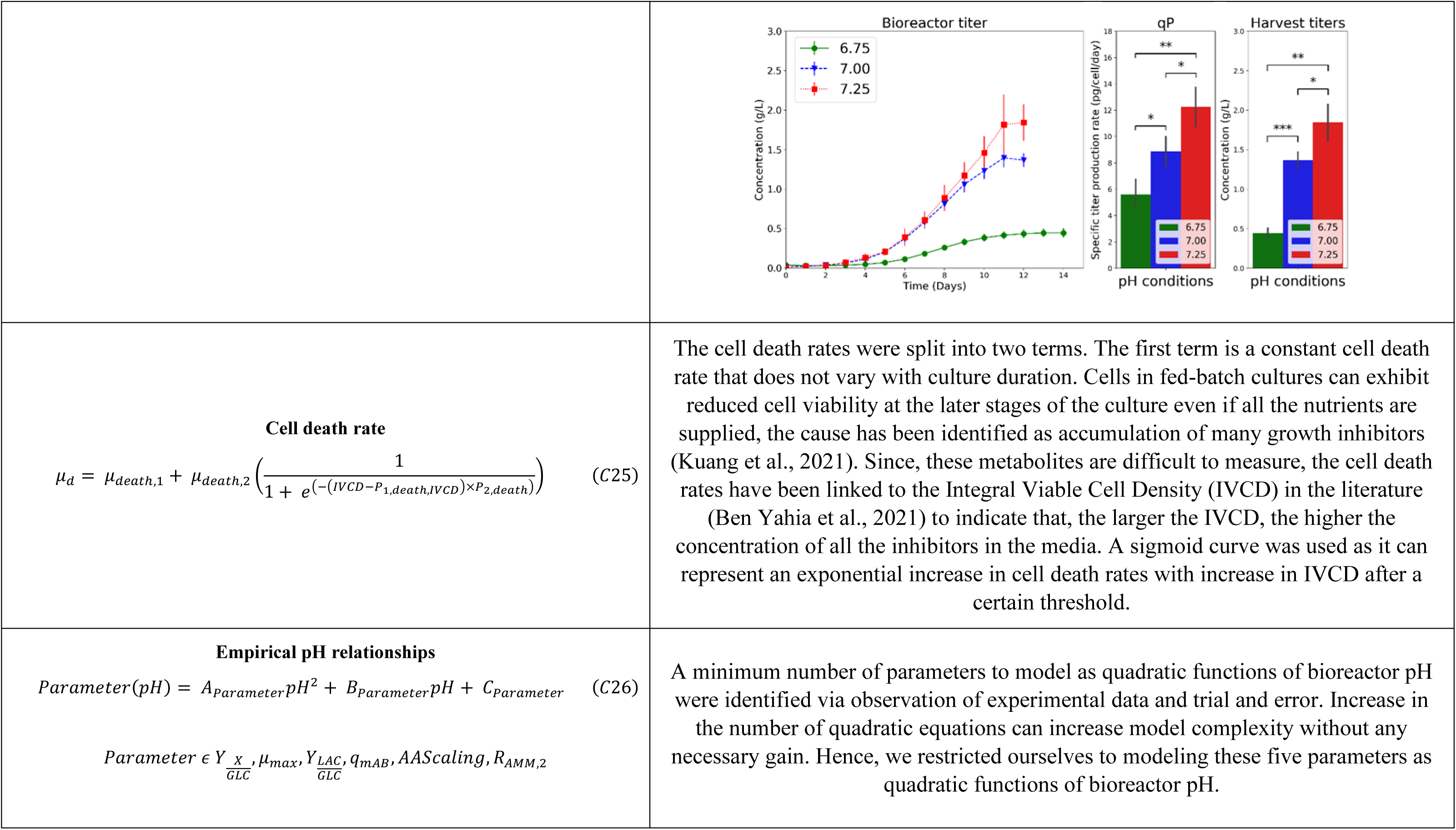
Kinetic equations. The kinetic equations in the model are semi-empirical and formulated based on the observed metabolic phenotypes in our previous work (Reddy et al., 2025).

### 10.1 Fed-Batch and Intensified Fed-Batch Mass Balances

**Figure C1:**
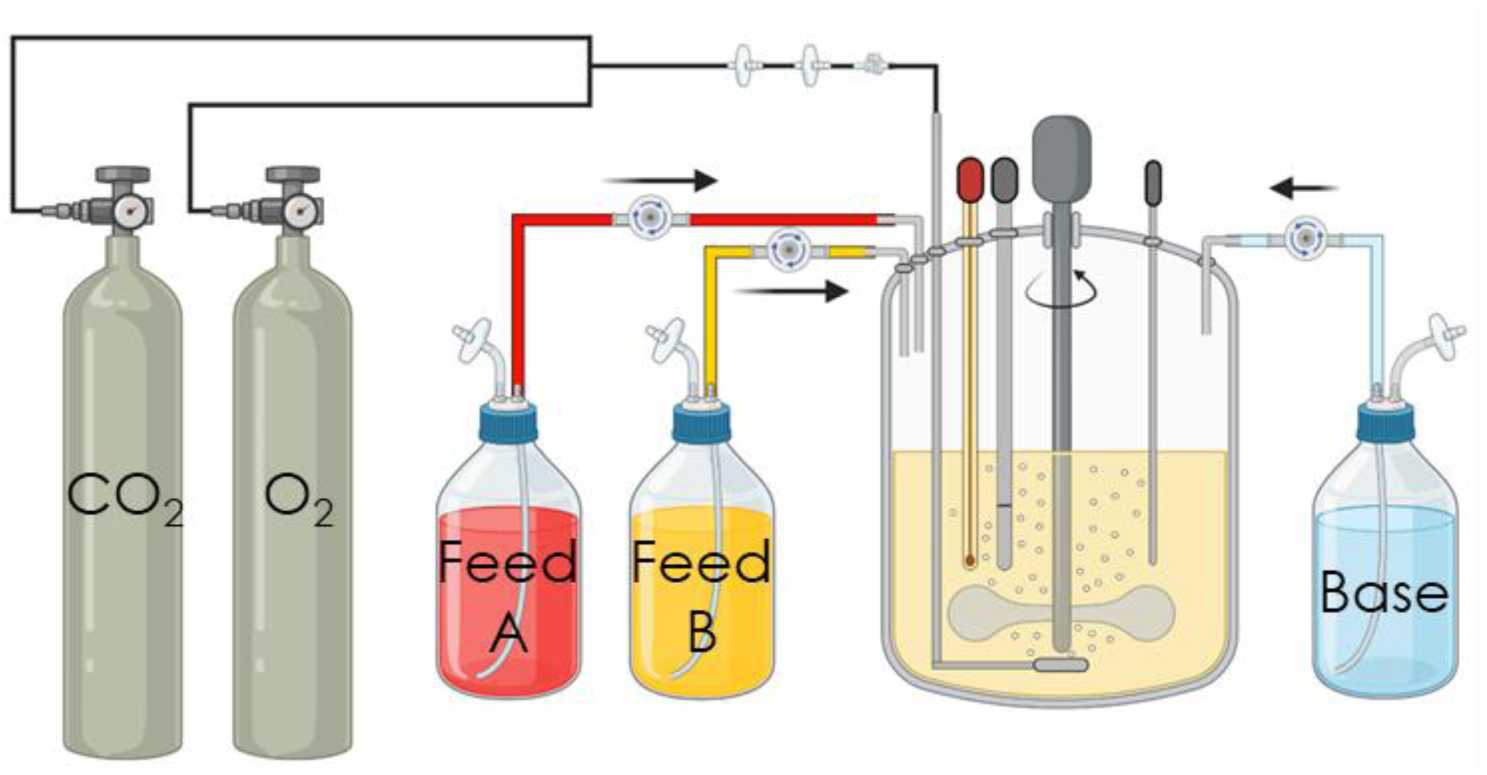
Fed-batch and intensified fed-batch bioreactor configuration.

The time-dependent fed-batch and intensified fed-batch mass balances are summarized in Table C2. Mass balances are performed on the bioreactor working volume, 𝑉_𝑅_, the viable cell density, [𝑉𝐶𝐷], all measured substrates which include glucose and amino acids, [𝑆], and all measured metabolites which include lactate, ammonia, and mAb, [𝑀]. For glutamine and ammonia specifically, the contribution of the glutamine degradation rate, 𝑘_𝑑,𝐺𝐿𝑁_, is considered in the mass balances as shown in Equations (C30 and C32). Depending on the culture conditions, ammonia may be produced as a metabolite or consumed as a substrate.

**Table C2:**
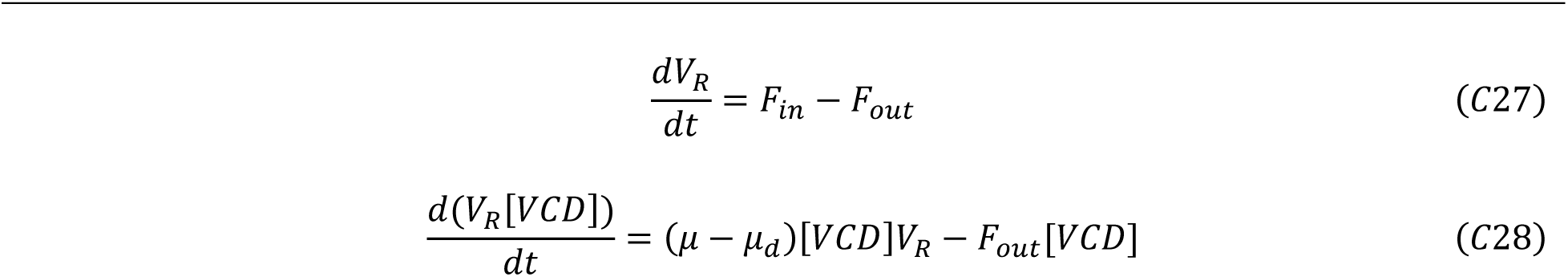
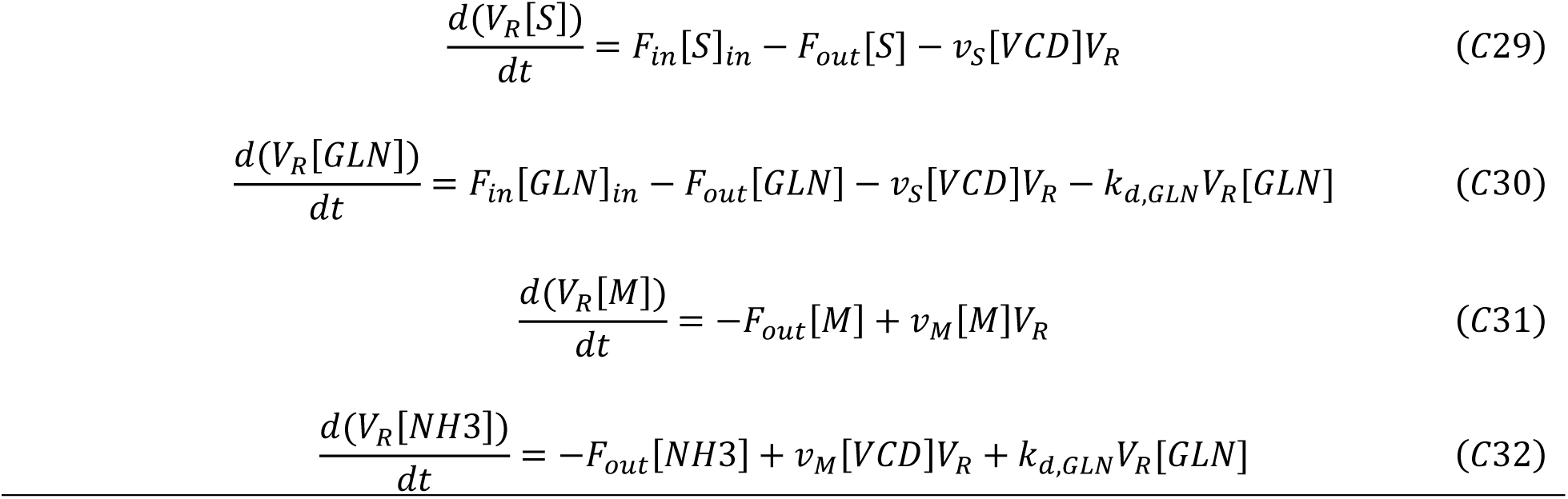
Time-dependent fed-batch and intensified fed-batch mass balances on the bioreactor working volume, the viable cell density, substrates, and metabolites.

𝜇 is the specific growth rate, 𝜇_𝑑_ is the specific death rate, 𝐹_𝑖𝑛_ is the total volumetric flowrate of feed introduced into the bioreactor, 𝐹_𝑜𝑢𝑡_ is the total volumetric flowrate of culture removed from the bioreactor, 𝑣_𝑠_ is the uptake flux of the substrate, and 𝑣_𝑀_ is the production flux of the metabolite.

The derivate term in the bioreactor mass balances is linearized according to the finite forward difference method. The balances are rearranged to project the bioreactor working volume, 𝑉_𝑅,𝑡+1_, viable cell density, [𝑉𝐶𝐷]_𝑡+1_, the extracellular substrate concentrations, [𝑆]_𝑡+1_, and extracellular metabolite concentrations, [𝑀]_𝑡+1_, at the next timepoint of the culture, 𝑡 + 1.

**Table C3:**
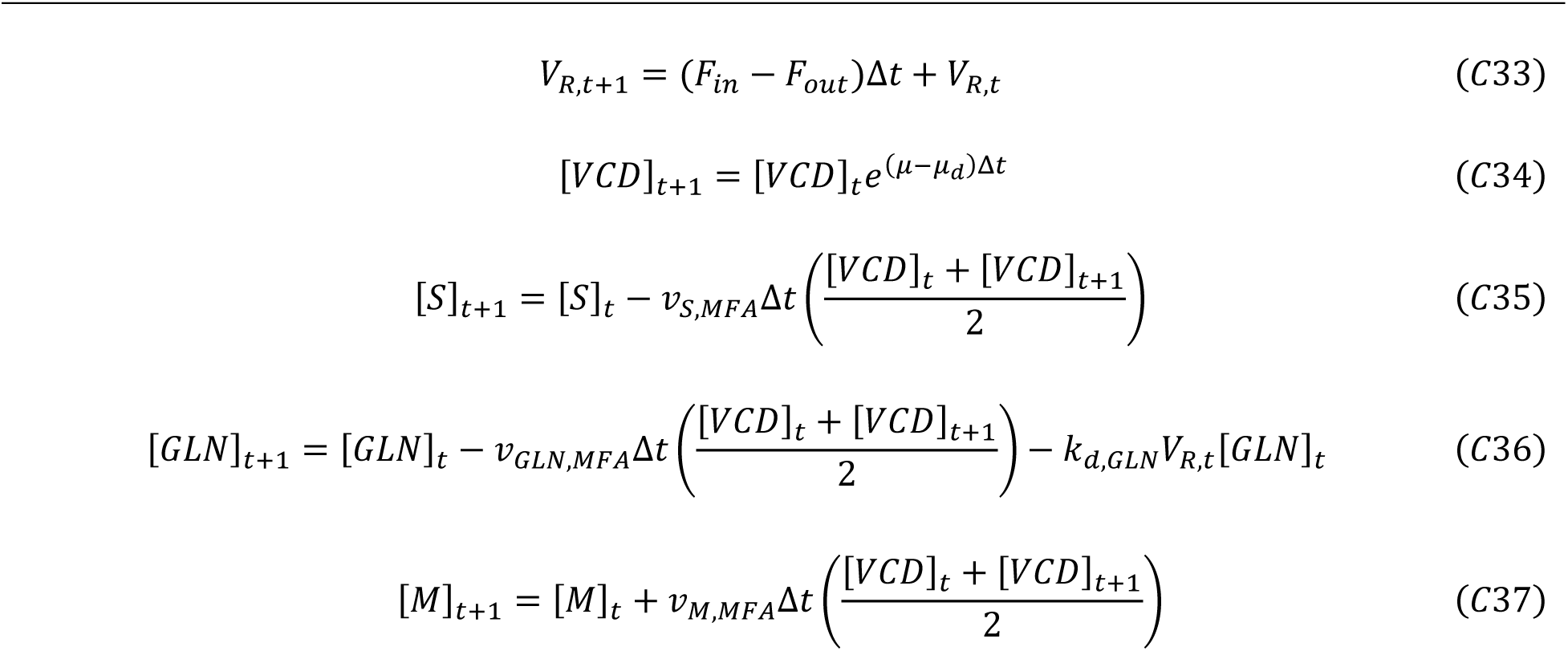
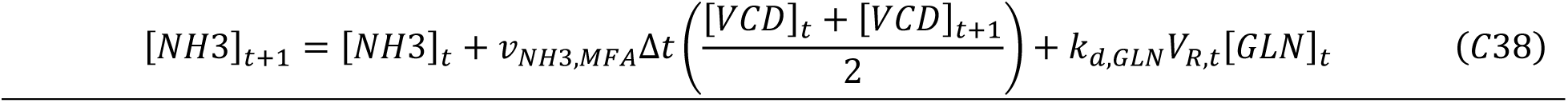
Linearized fed-batch and intensified fed-batch mass balances.

Δ𝑡 is the time step taken here as 0.1 days, 𝑣_𝑆,𝑀𝐹𝐴_ is the uptake flux of the substrate obtained from the MFA solution to the stoichiometric network, and 𝑣_𝑀,𝑀𝐹𝐴_ is the production flux of the metabolite obtained from the MFA solution to the stoichiometric network.

### 10.2 Perfusion Bioreactor Mass Balances

**Figure C2:**
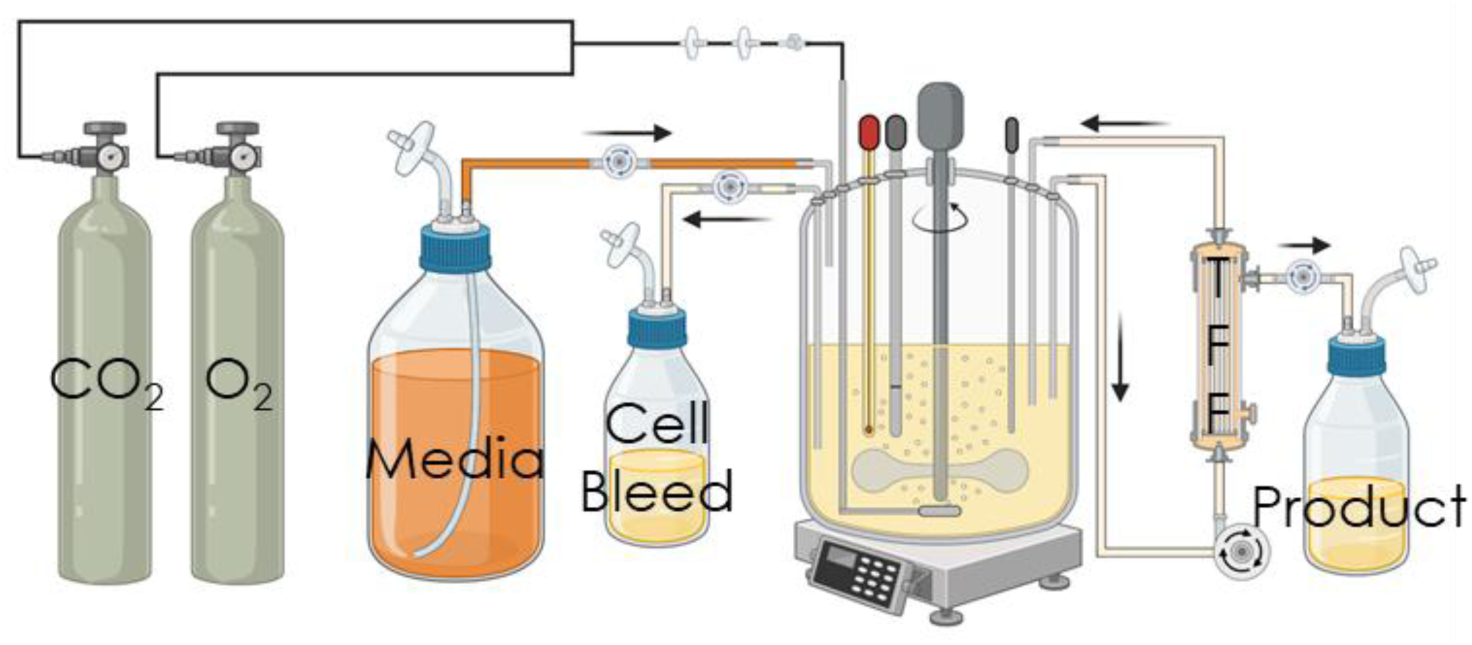
Perfusion bioreactor configuration.

The time-dependent perfusion mass balances are summarized in Table C3.

**Table C3:**
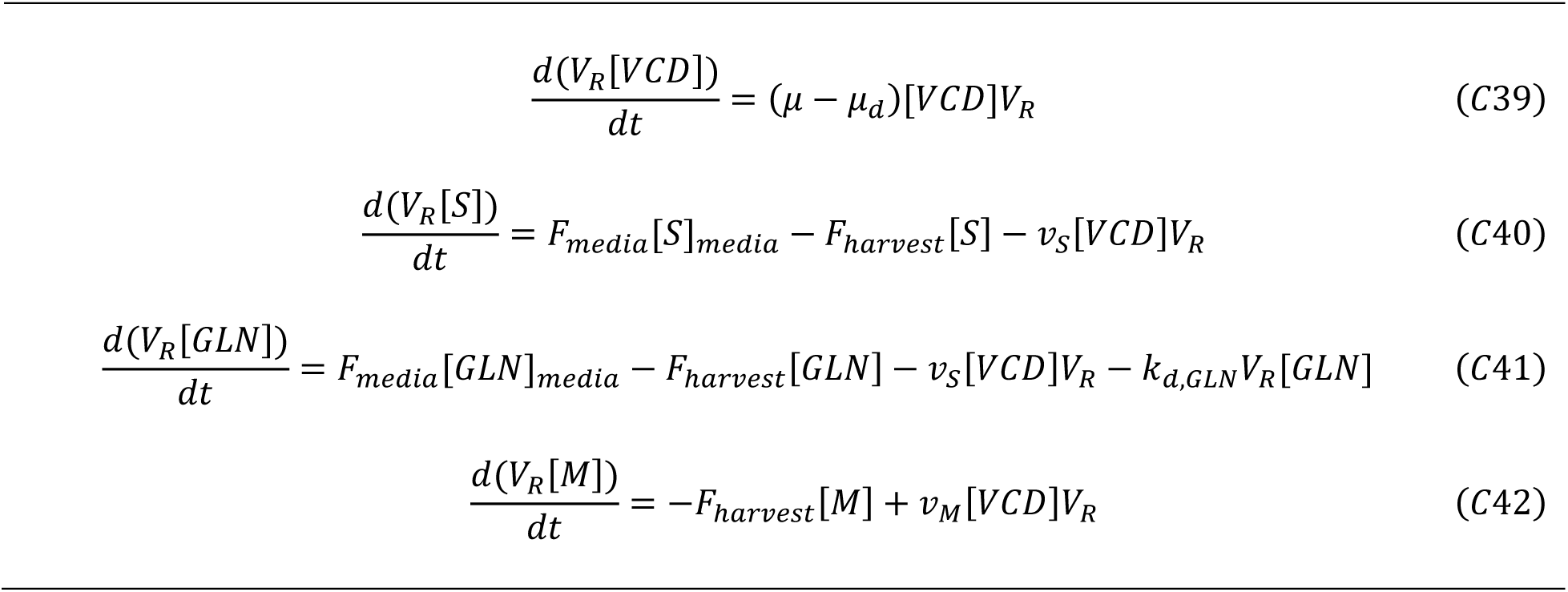
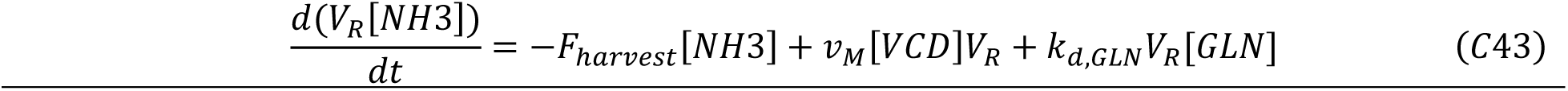
Time-dependent perfusion bioreactor mass balances on the viable cell density, substrates, and metabolites.

𝐹_𝑚𝑒𝑑𝑖𝑎_ is the volumetric flowrate of fresh perfusion media, [𝑆]_𝑚𝑒𝑑𝑖𝑎_ is the substrate concentration in the perfusion media, and 𝐹_ℎ𝑎𝑟𝑣𝑒𝑟𝑠𝑡_ is the volumetric flowrate of the harvest stream.

The perfusion bioreactor is operated with an intermittent cell bleed, hence the cell bleed is not included in the VCD mass balance of Equation (C39) and is identical to the fed-batch mass balance of Equation (C28). The cell bleed is engaged to control the specific growth rate by removing excess cells to enable steady state operation. The required daily bleed volume, 𝑉_𝑏𝑙𝑒𝑒𝑑_, removed during the intermittent cell bleed to maintain the VCD about the setpoint, [𝑉𝐶𝐷]_𝑠𝑝_, is calculated from a mass balance on the viable cells, Equation (C45).

After the bleed is performed, the bleed volume is replaced with fresh perfusion media to restore the bioreactor working volume. The extracellular substrate and metabolite concentrations after the cell bleed, [𝑆]_𝑝𝑜𝑠𝑡−𝑏𝑙𝑒𝑒𝑑_and [𝑀]_𝑝𝑜𝑠𝑡−𝑏𝑙𝑒𝑒𝑑_respectively, are updated accordingly in Equations (C50 and C53). Therefore, the perfusion mass balances describe substrate and metabolite concentrations in between and immediately after the intermittent cell bleeds.

**Table C4:**
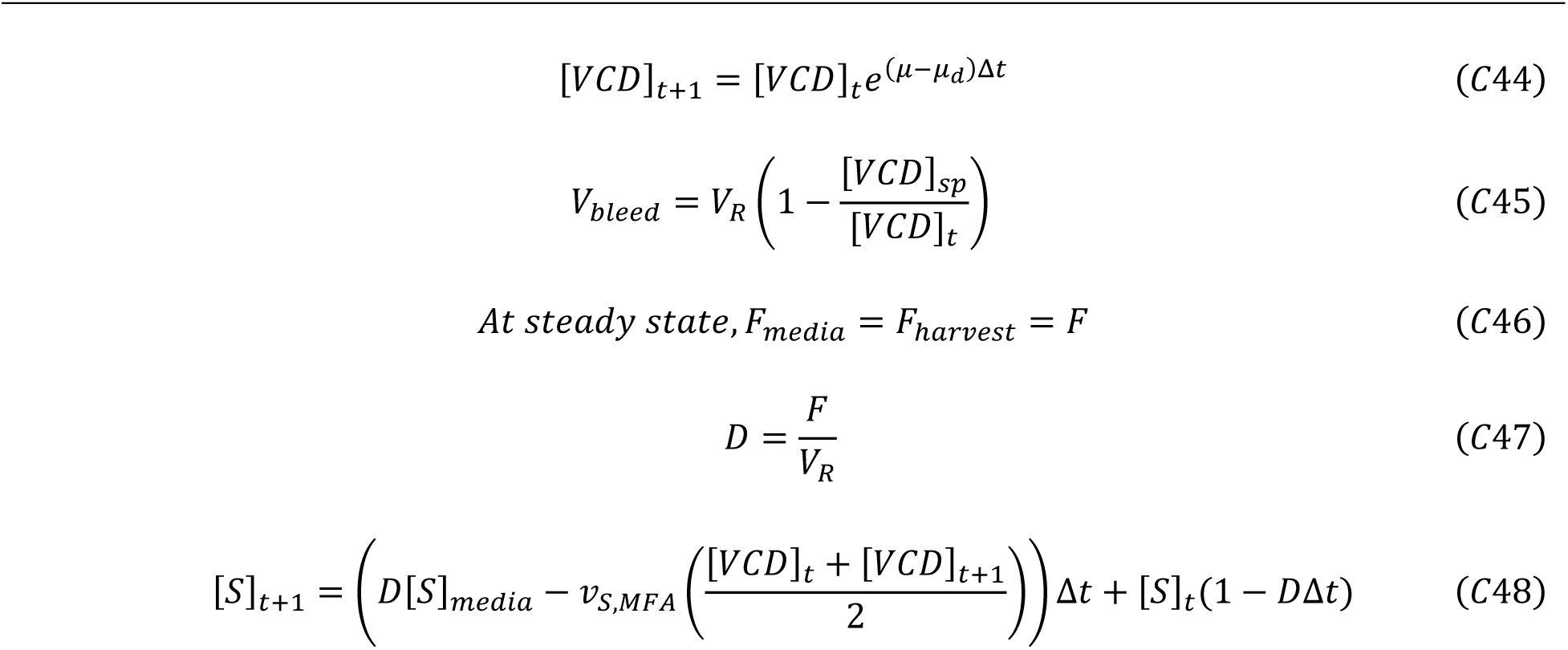
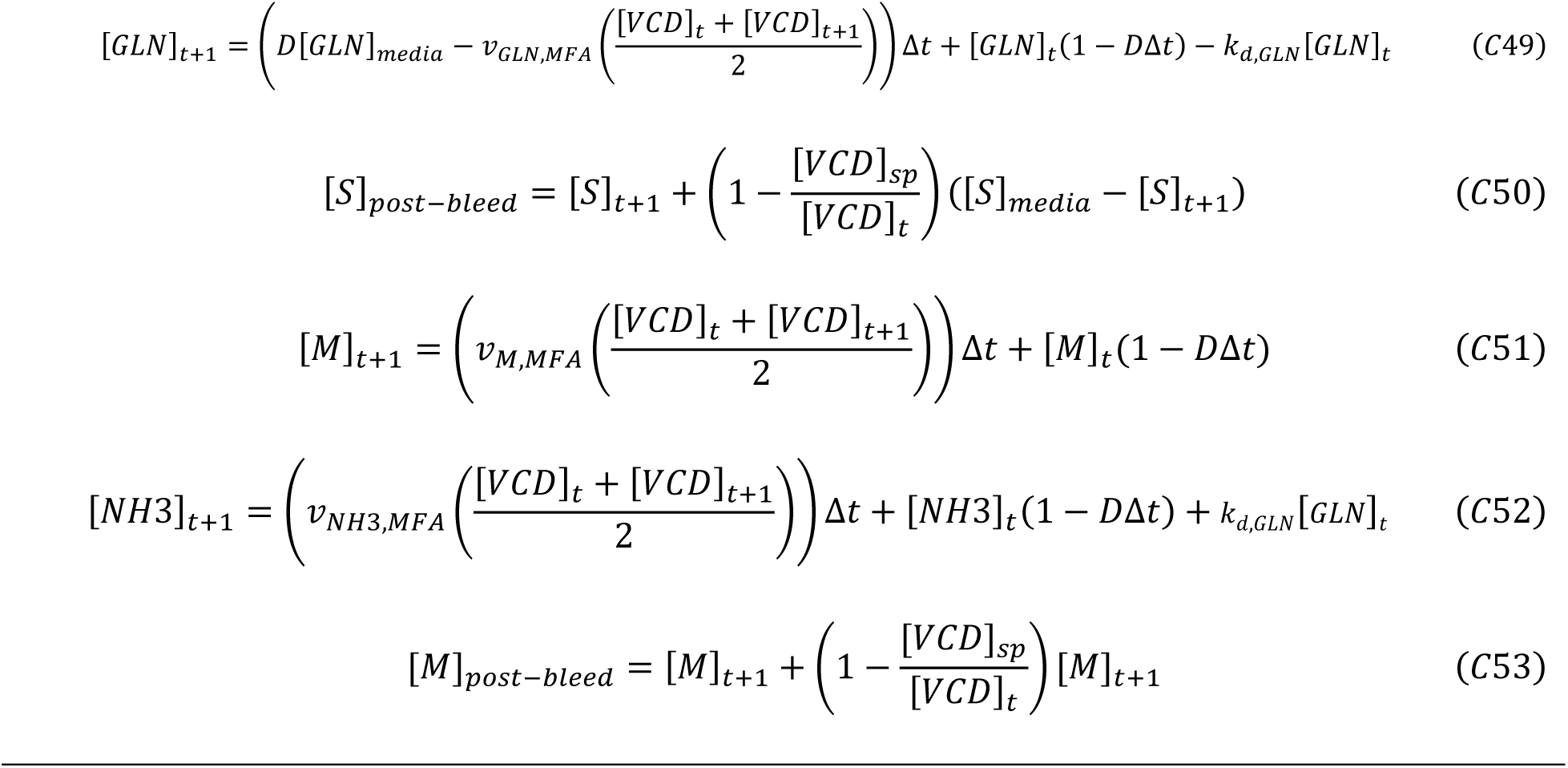
Linearized perfusion bioreactor mass balances.

At steady state, 𝐹_𝑚𝑒𝑑𝑖𝑎_ and 𝐹_ℎ𝑎𝑟𝑣𝑒𝑠𝑡_ are equivalent. 𝐷 is the perfusion rate.

## Notes

### Competing Interest Statement

The authors have declared no competing interest.

