## Supplementary Section for "A dynamic metabolic flux analysis (DMFA) model for performance predictions of diverse CHO cell culture process modes and conditions"

### S1 Biomass reaction calculations:

The amino acid composition of CHO-K1 cells have also been taken from the literature (Széliová et al., 2020). However, the composition of aspartate + asparagine and glutamate + glutamine in the protein content of CHO-k1 cells was reported. The ratio of aspartate to asparagine and glutamate to glutamine compositions were not reported. These ratios were taken from another literature study (Zamorano et al., 2010). Based on the following composition the molecular weight of the protein is 129 g/mol.

| Amino acid | Mole % in CHO biomass protein |
| --- | --- |
| Glycine | 8.6 |
| Leucine | 8.3 |
| Alanine | 7.7 |
| Lysine | 7.2 |
| Serine | 6.5 |
| Arginine | 6.3 |
| Proline | 5.5 |
| Valine | 5.0 |
| Threonine | 5.2 |
| Isoleucine | 3.8 |
| Phenylalanine | 3.4 |
| Tyrosine | 2.8 |
| Cysteine | 2.0 |
| Histidine | 2.0 |
| Methionine | 1.8 |
| Tryptophan | 1.0 |
| Asparagine | 4.6 |
| Aspartate | 5.6 |
| Glutamine | 5.1 |
| Glutamate | 7.6 |

This leads to the protein formation reaction

*0.086 Gly + 0.083 Leu + 0.077 Ala + 0.072 Lys + 0.065 Ser + 0.063 Arg + 0.055 Pro + 0.050 Val + 0.052 Thr + 0.038 Ile + 0.034 Phe + 0.028 Tyr + 0.020 Cys + 0.020 His + 0.018 Met + 0.010 Trp + 0.046 Asn + 0.056 Asp + 0.051 Gln + 0.076 Glu + 4 ATP -> 1 Protein*

The lipid composition for CHO cells was also taken from the literature (Zamorano et al., 2010). Based on the following composition the molecular weight of lipids is 676 g/mol.

| Lipid | Mole % per total lipid |
|---|---|
| PE (phosphatidylethanolamine) | 20 |
| PC (phosphatidylcholine) | 50 |
| SM (sphingomyelin) | 7 |
| PS (phosphatidylserine) | 7 |
| ST (cholesterol) | 16 |

This leads to the following reaction.

*0.155 PE + 0.358 PC + 0.038 PS + 0.263 SM + + 0.186 Cholesterol -> 1 total lipids*

Relative biomass composition for CHO-K1 cells has been published in the literature (Hagrot et al., 2019; Széliová et al., 2020). The average weight composition proteins, lipids, RNA, DNA, carbohydrates, and ash content per cell from three CHO-K1 cell lines that are used to calculate the biomass composition are listed below.

| Macromolecules | Composition (% weight per cell) |
|---|---|
| Protein | 53.0 |
| Lipid | 12.3 |
| DNA | 3.2 |
| RNA | 8.0 |
| Total carbohydrates | 1.6 |
| Remaining cellular material | 21.2 |

The elemental composition of CHO cell biomass is taken from the literature (Berrios et al., 2011).

| Element | Stoichiometry |
|---|---|
| C | 1 |
| H | 1.78 |
| O | 0.44 |
| N | 0.24 |

The molecular weight of C-mol of CHO cell biomass is 24.18 g/mol.

Based on the % composition of macromolecules, molecular weight of macromolecules, and the molecular weight of CHO cell biomass, the following overall biomass reaction is summarized.

*0.0993 Protein + 0.0047 Lipid + 0.0024 DNA + 0.0057 RNA + 0.0024 Total carbohydrates -> 1 Biomass*

The reactions leading to synthesis of DNA, RNA, Lipids, and total carbohydrates are taken from the literature (Zamorano et al., 2010). These reactions are lumped together to get the overall biomass reaction.

1 G6P + 8 ATP + 2 ASP + 1 GLY + 2 GLN -> 2 NADPH + 2 FUM + 2 GLU + 1 ATPRN

1 G6P + 8 ATP + 1 ASP + 1 GLY + 3 GLN -> 1 FUM + 3 GLU + 2 NADPH + 1 NADH + 1 GTPRN

1 G6P + 2 ATP -> 2 CO2 + 2 NADPH + 1 NADH + 1 UTPRN

1 GLN + 1 G6P + 2 ATP -> 2 CO2 + 2 NADPH + 1 NADH + 1 GLU + 1 CTPRN

0.28 UTPRN + 0.28 ATPRN + 0.22 GTPRN + 0.22 CTPRN -> 1 RNA

1 ATPRN -> 1 dATP

1 GTPRN -> 1 dGTP

1 UTPRN -> 1 dTTP

1 CTPRN -> 1 dCTP

0.28 dATP + 0.22 dGTP + 0.28 dTTP + 0.22 dCTP -> 1 DNA

0.5 G6P + 23.5 ATP + 34 NADH + 18 AcCoA + 1 Ethn -> 1 PE

0.5 G6P + 23.5 ATP + 34 NADH + 18 AcCoA + 1 Cho -> 1 PC

1 SER + 0.5 G6P + 23.5 ATP + 34 NADH + 18 AcCoA -> 1 PS

16 AcCoA + 1 SER + 1 Cho + 16 ATP + 29 NADPH -> 1 SM + 2 CO2

18 AcCoA + 18 ATP + 14 NADPH -> 1 Cholesterol + 6 CO2

0.155 PE + 0.358 PC + 0.038 PS + 0.263 SM + + 0.186 Cholesterol -> 1 ME

AAs + 4 ATP -> 1 Protein

1 G6P + 1 ATP -> 1 TC

The above reactions are lumped together to yield the following biomass reaction.

0.0077 ALA + 0.0063 ARG + 0.0046 ASN + 0.0119 ASP + 0.0020 CYS + 0.0167 GLN + 0.0126 GLY + 0.0020 HIS + 0.0038 ILE + 0.0082 LEU + 0.0072 LYS + 0.0018 MET + 0.0034 PHE + 0.0071 SER + 0.0052 THR + 0.001 TRP + 0.0028 TYR + 0.0050 VAL + 0.5444 ATP +

0.0025 $NH_3$ + 0.1178 NADH + 0.0843 AcCoA + 0.0123 G6P -> 0.0132 $CO_2$ + 0.0041 GLU + 0.0063 FUM + 1.0 Biomass

The dry weight of CHO-K1 cells is taken from nine different measurements of CHO-K1 dry cell weight in the literature (Széliová et al., 2020), the value is 247 pg/cell.

The growth rate is converted to the biomass flux (pmol biomass/cell/day) using the following reaction.

$$q_{biom} = \mu \frac{dry\ weight\ per\ cell \times (1 - ash\ fraction)}{Molecular\ weight\ of\ biomass}$$

### Figures

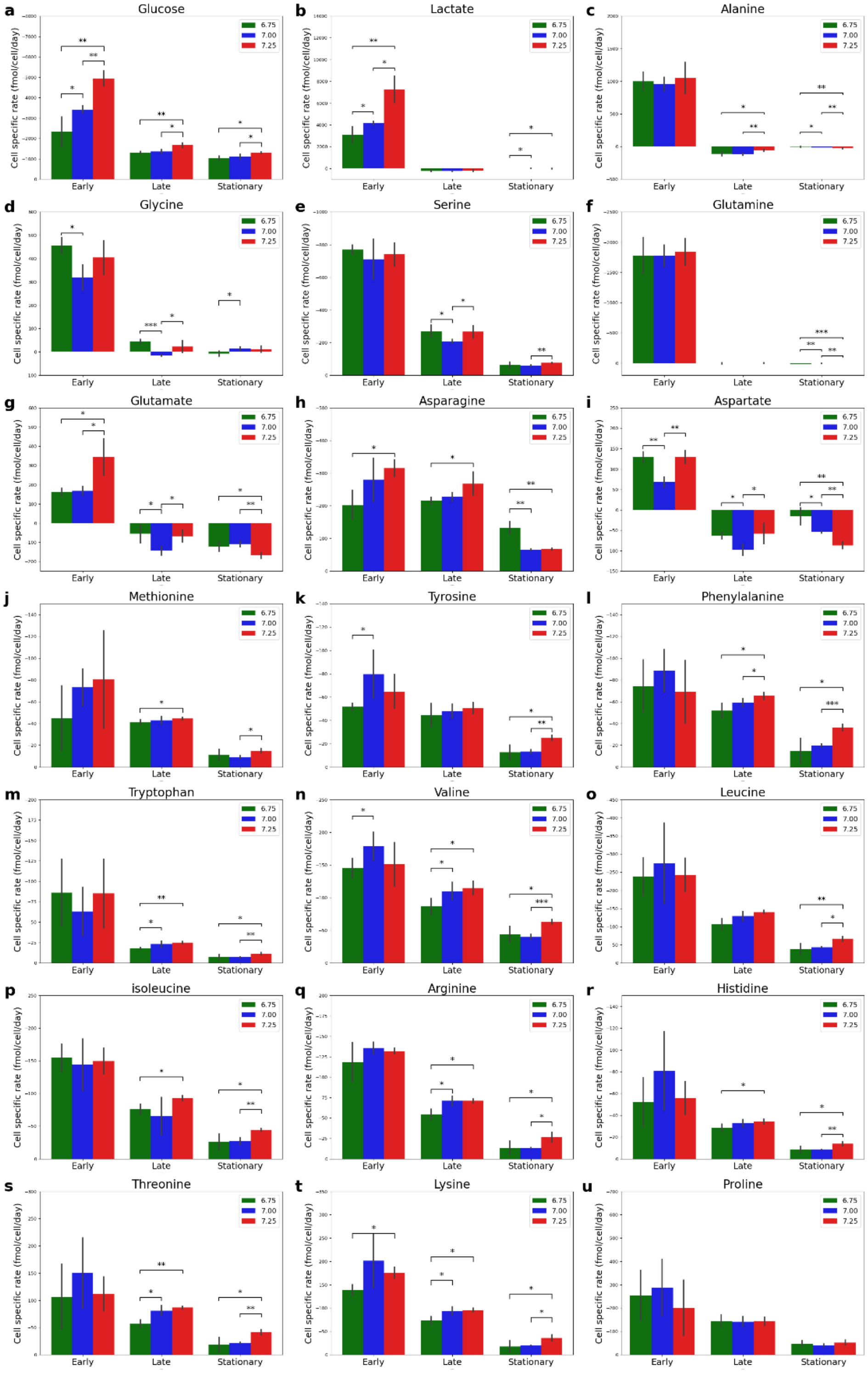


**Figure S1:** Uptake and secretion rates of measured metabolites under various bioreactor pH conditions at three culture phases.

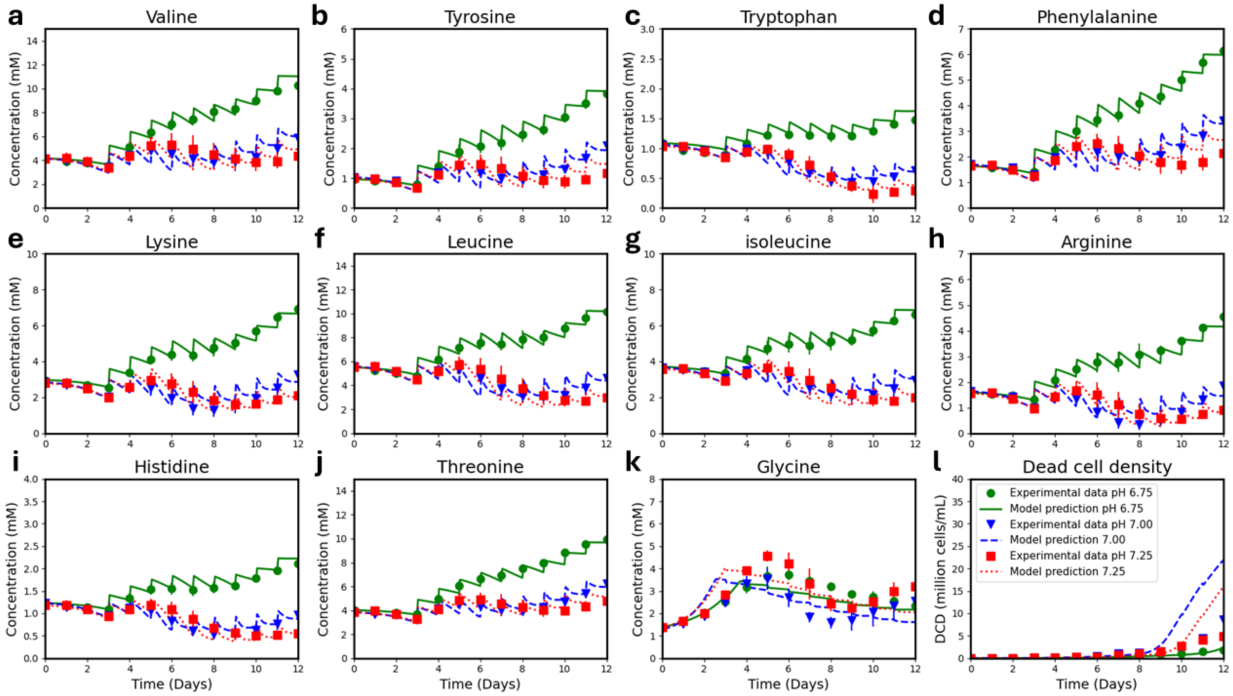


**Figure S2**: Predictions of impact of bioreactor pH on metabolite concentrations

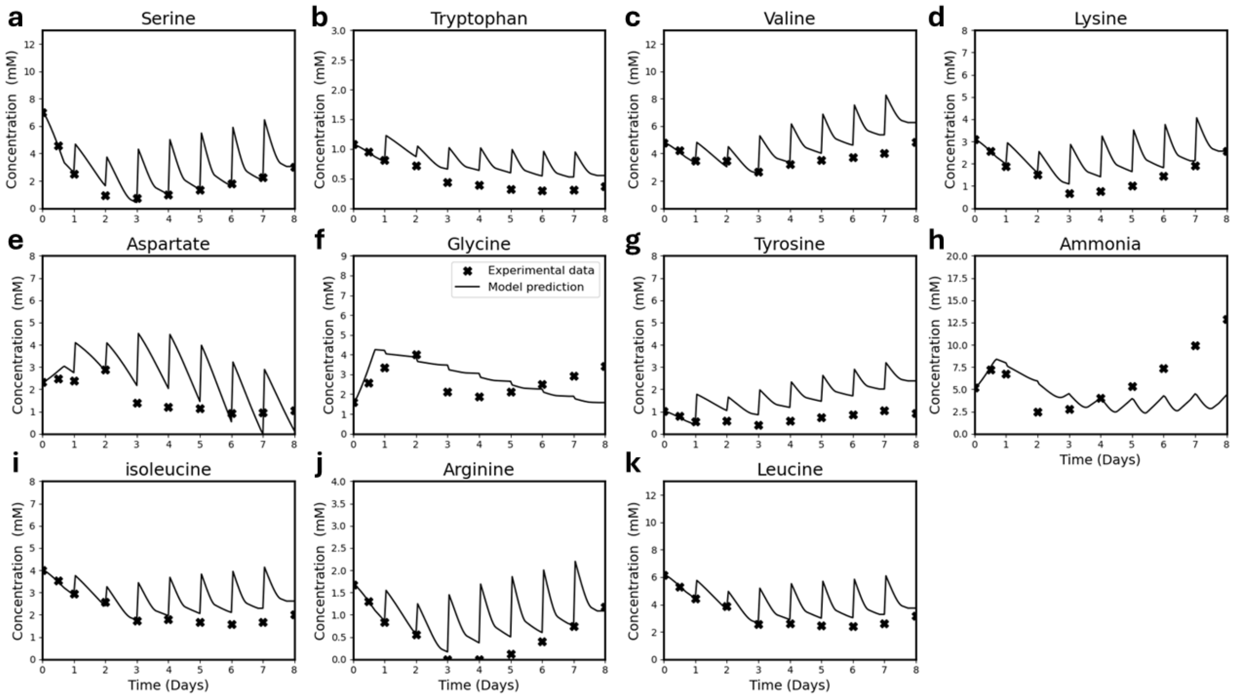


**Figure S3:** Predictions of intensified fed-batch culture performance

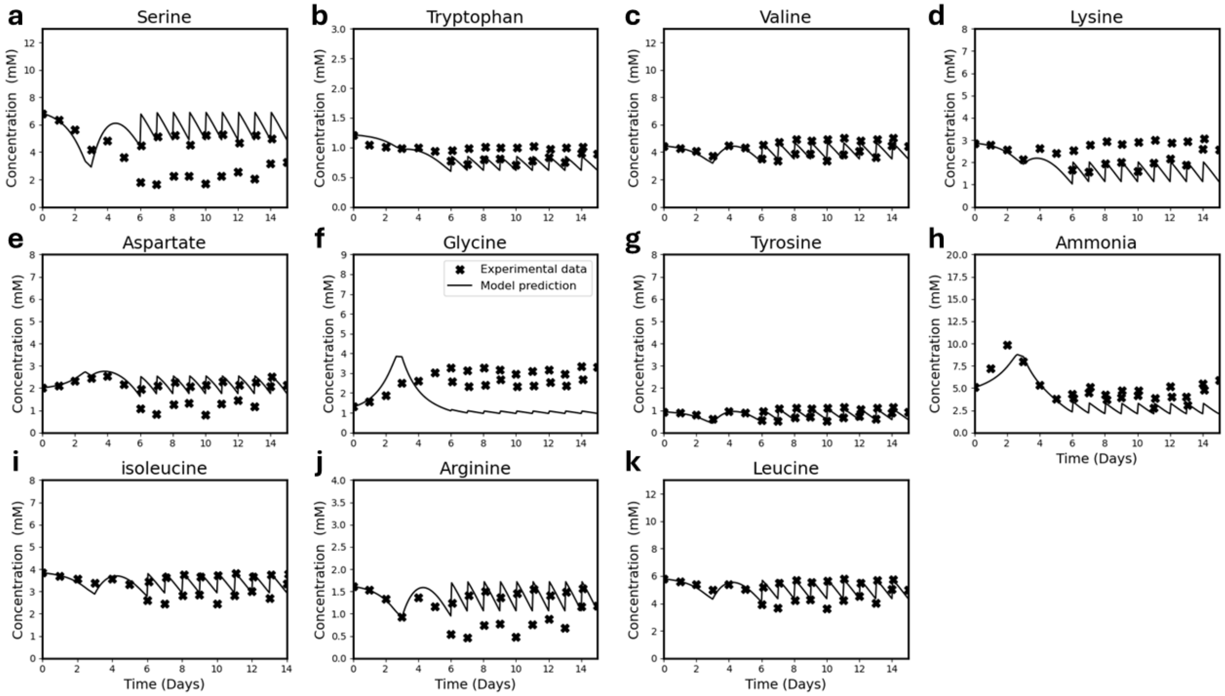


**Figure S4:** Predictions of perfusion bioreactor process performance

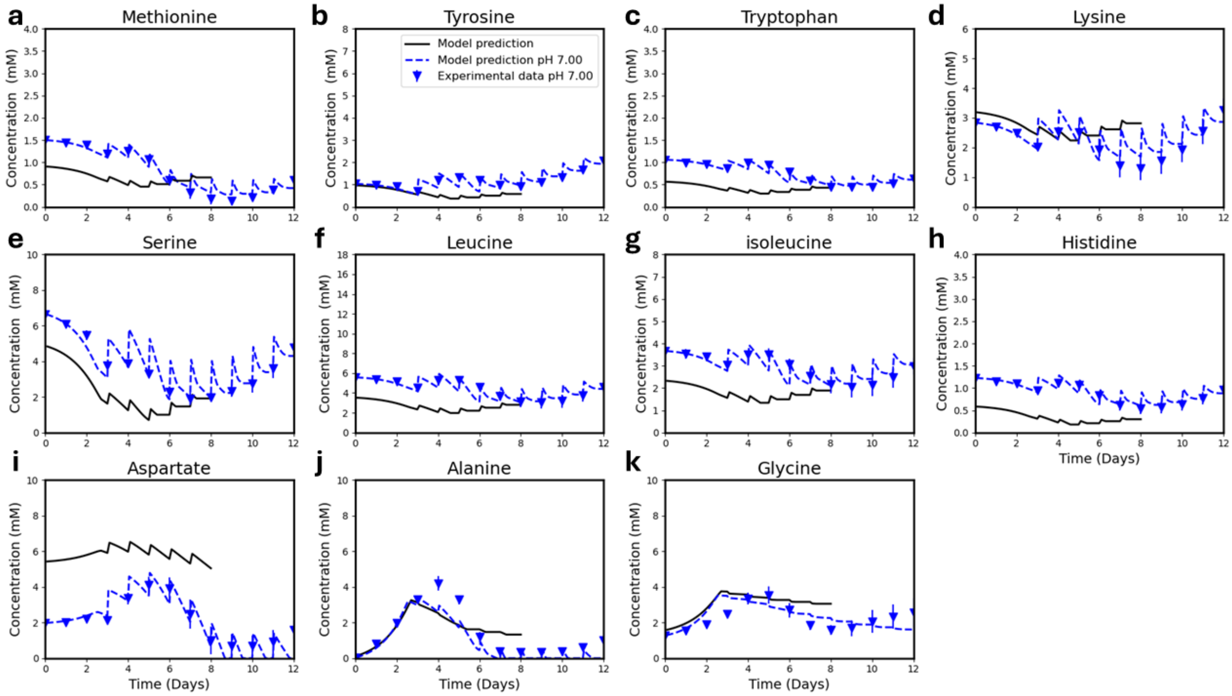


**Figure S5:** Predictions of AMBIC reference basal and feed media performance.
